# Dynamic arrest and aging of biomolecular condensates are regulated by low-complexity domains, RNA and biochemical activity

**DOI:** 10.1101/2021.02.26.433003

**Authors:** Miriam Linsenmeier, Maria Hondele, Fulvio Grigolato, Eleonora Secchi, Karsten Weis, Paolo Arosio

## Abstract

Biomolecular condensates require suitable material properties to properly carry out their function. Yet, modulators that affect the material properties of condensates have remained largely unexplored.

Here we apply Differential Dynamic Microscopy (DDM) to probe the material properties of an *in vitro* model of processing bodies (P-bodies) consisting of condensates formed by the DEAD-box ATPase Dhh1 in the presence of ATP and RNA. DDM allows us to measure the viscosity of liquid droplets and to distinguish between liquid-like and gel- or glass-like condensates. By applying this single-droplet technique we show that condensates within the same population exhibit a distribution of material properties, which can be drastically affected by several modulators such as the low-complexity domains (LCDs) of the protein, the protein/RNA ratio, the type of RNA as well as the enzymatic activity.

In particular, we show that structured RNA leads to a large fraction of dynamically arrested condensates with respect to unstructured polyuridylic acid (polyU), emphasizing the role of RNA structure in condensate dynamics. We further demonstrate that the ageing of the condensates and the formation of gel or glass-like structures is reduced by promoting the enzymatic ATPase activity of Dhh1 and the rate of droplet formation and dissolution.

Our study shows that not only the reversible formation and dissolution of condensates but also their material properties are regulated on several levels, and that biochemical activity and material turnover can be important to maintain fluid-like properties over time.

---

The ability of cells to form compartments is crucial to coordinate a variety of reactions in space and time. In addition to membrane-bound compartments, it is becoming increasingly clear that cells form membrane-less organelles by liquid-liquid phase separation (LLPS) of proteins and nucleic acids (Aguzzi and Altmeyer, 2016; Banani et al., 2017; Brangwynne et al., 2009, 2015). The dynamic formation and dissolution of these biomolecular condensates is governed by a variety of intermolecular interactions (Wheeler and Hyman, 2018), which involve multivalency and repetitive sequence patterns (Bouchard et al., 2018; Harmon et al., 2017; Li et al., 2012; Mittag and Parker, 2018). These multivalent interactions can be promoted by intrinsically disordered protein sequences known as low complexity domains (LCDs) (Hughes et al., 2018), by globular protein-protein interactions (Bouchard et al., 2018) or by RNA-protein interactions (Lin et al., 2015). An important feature of phase separating systems is the responsiveness to factors such as changes in ionic strength and pH, but also factors like ATP (Mugler et al., 2016; Patel et al., 2017), nucleic acids (Begovic et al., 2020; Du and Chen, 2018; Elbaum-Garfinkle et al., 2015; Langdon et al., 2018; Nott et al., 2015; Sachdev et al., 2019; Tauber et al., 2020) and small molecules (Wheeler et al., 2019).

While a lot of attention has been dedicated to the effect of different factors on the formation and dissolution of biomolecular condensates, mechanisms that control their material properties have remained much less explored. Yet, suitable material properties (viscosity, elasticity, surface tension) are likely crucial for the proper physiological function of biomolecular condensates and misregulation of these properties may lead to pathologies (Mathieu et al., 2020).

Biological condensates contain molecular networks whose formation is mediated by multivalent interactions (Harmon et al., 2017; Posey et al., 2018) and can therefore be considered as structured network fluids (Guillén-Boixet et al., 2020). A variety of material properties ranging from liquid-like to dynamically arrested gel- or glass-like have been reported. In some cases, maturation from a liquid-like state into such arrested states has been observed over time (Woodruff et al., 2017; Zeng et al., 2018), potentially leading to the formation of aberrant protein aggregates or amyloids. This pathological liquid-to-solid phase transition has been associated with neurodegenerative diseases (Babinchak et al., 2019; Molliex et al., 2015; Patel et al., 2015; Ray et al., 2020; Wegmann et al., 2018). Understanding the regulation of the materials properties after condensate formation and their evolution over time is particularly important. For instance, for condensates hosting biochemical reactions, fluidity is typically required to recruit client molecules and rapidly release products after processing (Hondele et al., 2019). By contrast, other condensates may require a certain level of rigidity to form a stable structural matrix (Boke et al., 2016; Dufresne et al., 2009; Hubstenberger et al., 2013; Putnam et al., 2019; Woodruff et al., 2017). The assessment of the material properties of condensates, especially of dynamically arrested states, is still limited both *in vivo* and *in vitro*. In addition, the molecular factors that modulate these material properties have remained largely unraveled.

The distinction between liquid-like and gel-/glass-like materials requires a technique capable to probe the dynamics of the systems and cannot rely on structural information only (Janssen, 2018). In this context, several techniques have been developed in soft matter physics, including particle tracking and optical tweezers (Elbaum-Garfinkle et al., 2015; Jawerth et al., 2019, 2018). However, under some conditions, these techniques could be challenging to implement.

Here, for the first time, we apply Differential Dynamic Microscopy (DDM) to probe the material properties of *in vitro* models of biomolecular condensates. DDM probes the microscopic dynamics of the condensate by monitoring fluctuations in the intensity of scattered light over time. The technique is not invasive and can be applied in combination with nanoparticle tracers with size below the optical resolution. This allows to probe also small condensates that would exclude the large particle tracers that are conventionally required for particle tracking experiments (Cerbino and Trappe, 2008). A key advantage of the technique consists in the possibility to analyze individual condensates with sizes ranging from a few to hundreds of microns, therefore providing information on the distribution of material properties within a population of condensates. Moreover, DDM can be performed on a simple widefield microscope in bright field mode without the requirement of additional equipment. The technique provides an attractive opportunity to probe the material properties of condensates as a function of several molecular determinants and over time.

In this work, we apply DDM to investigate the material properties of condensates formed by the P-body-associated DEAD-box ATPase Dhh1 depending on LLPS-relevant factors such as ATP and RNA. This protein has several interesting features to explore the relationship between biochemical activity, dynamics of formation and dissolution of the condensates and their material properties.

Dhh1 has a globular core consisting of two RecA-like domains that contain the binding sites for RNA and ATP. These core domains are connected by a linker and are flanked by two low-complexity domains (LCDs) (Cheng et al., 2005; Mugler et al., 2016; Protter et al., 2018) (Fig. 1A).

**Figure 1.**
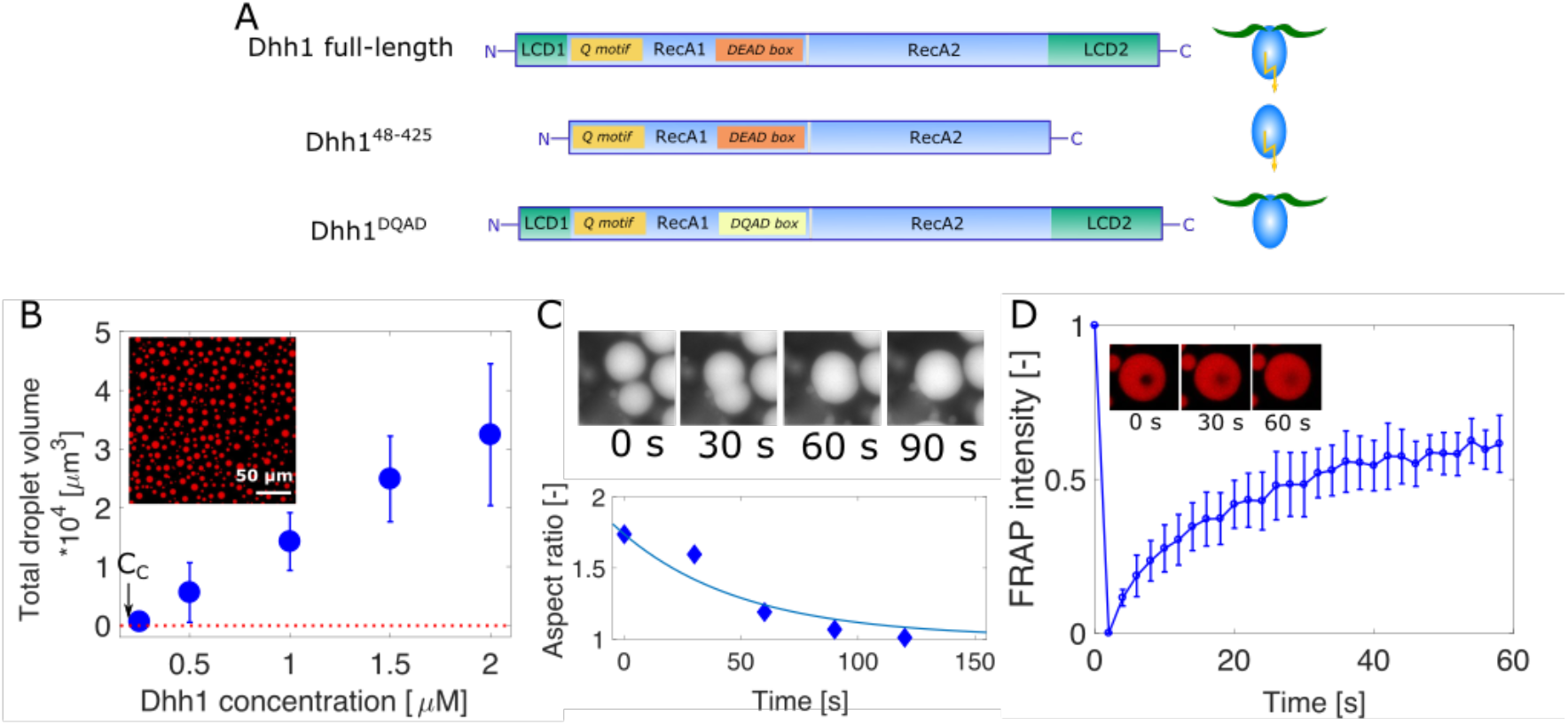
*In vitro* phase separation of Dhh1. **(A)** Dhh1 variants used in this study: full-length Dhh1, consisting of two RecA domains as core, flanked by an N- and a C-terminal low-complexity domain. Q motif and DEAD box are important motifs to bind and hydrolyze ATP. The Dhh1^48-425^ construct lacks both the N- and C-terminal LCD, while in the Dhh1^DQAD^ variant a single glutamate has been substituted with a glutamine, leading to an ATP hydrolysis-deficient variant. **(B)** Effect of Dhh1 concentration on droplet formation. Dhh1 solutions at increasing protein concentrations were supplied with 5 mM ATP / MgCl2 and 0.5 mg/ml polyU and imaged using widefield fluorescence microscopy providing a critical concentration (CC) equal to 0.2 µM. Error bars represent the standard deviation of droplets analyzed in three different samples. **(C)** Fusion and relaxation of two droplets into one single droplet, quantified by measuring the aspect ratio of two fusing droplets over time. **(D)** FRAP of the droplets shown in (B), showing a recovery of 61 ± 2 %.

While ATP binding and hydrolysis are mediated by the DEAD-box sequence and the Q motif (Linder and Jankowsky, 2011), the interaction with RNA is established in a sequence-independent manner *via* electrostatic interactions between the phosphate backbone of the RNA molecule and a positively charged cleft on the two RecA domains (Cheng et al., 2005). The binding of Dhh1 to ATP and RNA has been shown to be important to promote the formation of both P-bodies *in vivo* and for reconstituted liquid-like droplets *in vitro*. Furthermore, ATP hydrolysis by DEAD-box ATPases triggers RNA release from P-bodies and stress granules, and therefore offers an important possibility to regulate the disassembly of such bodies (Hondele et al., 2019; Mugler et al., 2016).

An attractive opportunity offered by this system is the possibility to investigate the effect of biochemical activity on the material properties of the condensates since there are several mechanisms that can modulate the intrinsic ATPase activity of Dhh1. For instance, the ATPase activity can be diminished by substituting a single amino acid in the DEAD-box of the protein, resulting in the exchange of a glutamate (DEAD) to a glutamine (DQAD). This ATP hydrolysis-deficient Dhh1^DQAD^ mutant forms constitutive processing bodies in yeast cells due to the impaired release of RNA from Dhh1, which underlies P-body dissolution. On the contrary, the ATPase activity can be stimulated by the P-body-associated factor Not1 (Mathys et al., 2014; Mugler et al., 2016), which leads to enhanced dynamics of P-bodies *in vivo* and to the dissolution of phase-separated droplets *in vitro*. In reconstituted systems, the hydrolyzed ATP can be regenerated, e.g., by using creatine kinase which transfers a phosphate residue on the released ADP molecule (using creatine phosphate as a donor) and the recycled ATP molecules can in turn re-promote phase separation in a cyclic way. Such a continuous, “fuel”-driven turnover (Tena-Solsona et al., 2018; Weber et al., 2019) of the droplet material keeps the phase separated system out of equilibrium and might ensure fluidity over time, preventing or delaying maturation to more solid-like, less dynamic states (Hondele et al., 2019).

Here we show that the material properties of Dhh1 condensates well as their maturation over time are controlled by intrinsic features encoded in the protein sequence (LCDs) as well as extrinsic factors (ATP hydrolysis, RNA). In particular, the lack of LCDs, the presence of structured RNA and the absence of enzymatic activity largely decreases the fluidity of the condensates, leading to their dynamic arrest.

Moreover, by applying DDM we show that under most of the investigated conditions populations of condensates from the same Dhh1 sample exhibit a distribution of material properties, including subpopulations of low viscous droplets, liquid-like droplets with high viscosities and dynamically arrested gel-/glass-like condensates.

Our results show that not only the formation of liquid-liquid phase separated condensates but also their material properties are carefully regulated on several levels.

## Results

### ATP, LCDs and RNA control the formation and dissolution of liquid-like protein-rich condensates

Condensates of mCherry-tagged full-length Dhh1 (Dhh1-mCh) were formed in an aqueous buffer of 90 mM KCl, 30 mM HEPES-KOH, pH 7.4 and 2 mM MgCl_2_ in the presence of 0.5 mg/ml of the RNA analog polyuridylic acid (polyU) and 5 mM ATP / MgCl_2_ (Fig. 1). Above the threshold solubility limit at a critical concentration of C_C_ = 0.2 µM, the droplet volume increased linearly with increasing mCh-Dhh1 concentration, highly suggestive of a phase transition (Fig. 1B). In absence of ATP and polyU, in the same buffer conditions, Dhh1 remained soluble up to around 500 µM (Suppl. Fig. 1). We observed fusion and relaxation of adjacent droplets into larger ones, indicative of a liquid-like character of these protein-rich condensates (Fig. 1C). These results are consistent with the results of our previous analysis performed in droplet microfluidics, which shows coalescence of Dhh1 condensates over time (Linsenmeier et al., 2019).

Analysis of the protein-rich droplets by fluorescence recovery after photobleaching (FRAP) showed an average recovery of the intensity to 61 ± 2% after 60 seconds (Fig. 1D).

We next investigated the effects of ATP, LCDs and polyU on the formation of the protein-rich condensates. To this aim, we analyzed an ATP-hydrolysis-deficient variant (Dhh1^DQAD^) as well as a truncated variant lacking the LCDs (Dhh1^48-425^), in addition to full-length Dhh1 (Fig. 1A). For all constructs, in the absence of ATP no LLPS was observed under the reference buffer conditions, even upon addition of only polyU (Suppl. Fig. S2, S3, S4). These results were confirmed by dynamic light scattering (DLS), which shows an average hydrodynamic diameter of 9.9 ± 3.0 nm for the full-length protein.

In contrast, the addition of an excess of 5 mM ATP / MgCl_2_ to a 5 µM Dhh1 solution resulted in the formation of non-spherical, protein particles with average hydrodynamic diameter of 605 ± 210 nm, as analyzed by a combination of dynamic light scattering and microscopy (Fig. 2A). This result, which was confirmed by the decrease in the amount of soluble monomer in presence of ATP measured by size exclusion chromatography coupled with UV absorbance (Fig. 2B), suggests an increase of intermolecular interactions upon addition of ATP. No such non-spherical particles were formed upon addition of GTP (even up to 25 mM, Suppl. Fig. 5A), excluding an unspecific effect of the ionic strength of the nucleotide. Similar results were observed for the catalytically inactive Dhh1^DQAD^ variant (Suppl. Fig. 5B-D) and the LCD-lacking Dhh1^48-425^ construct (Suppl. Fig. 5E-G).

**Figure 2.**
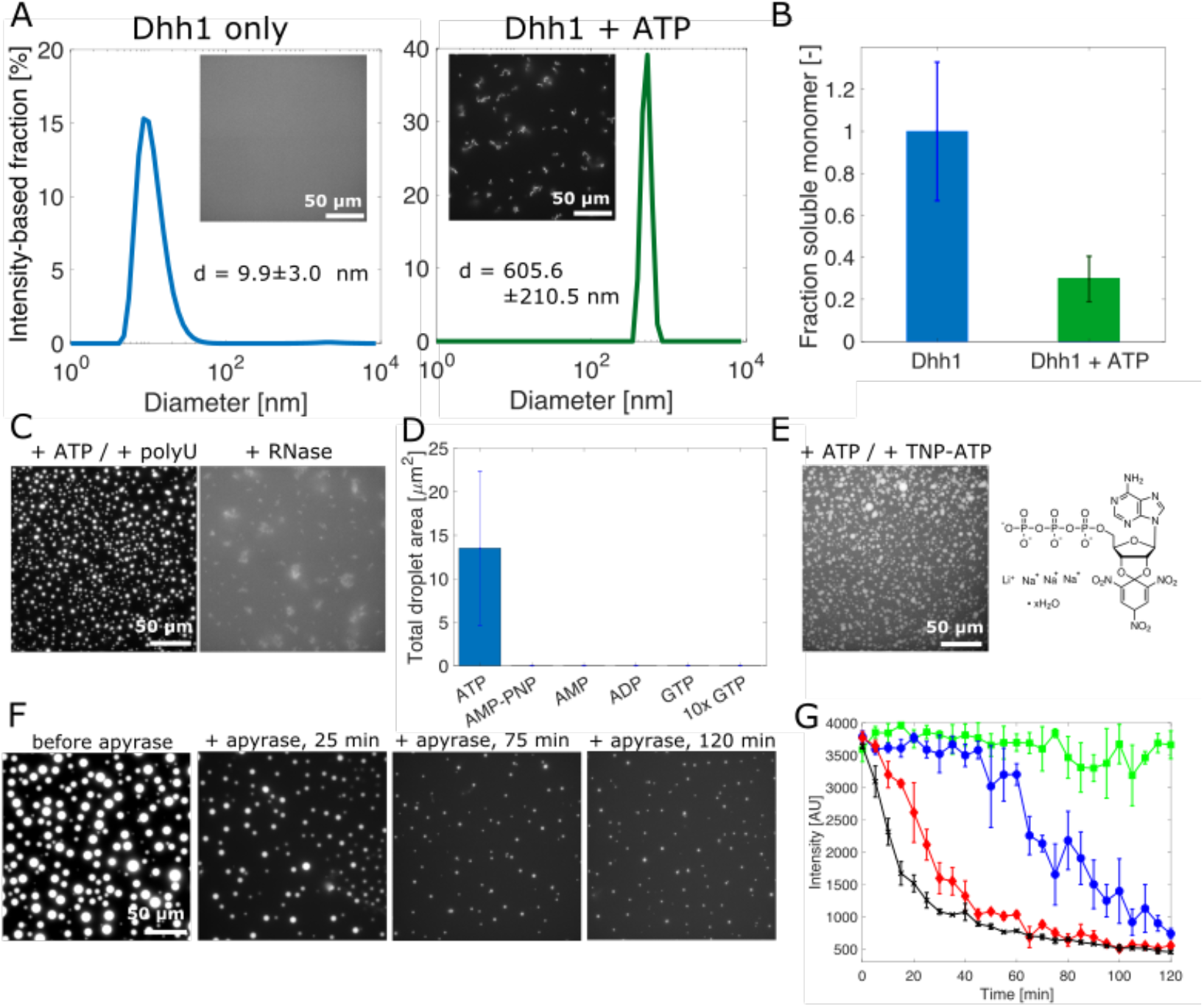
ATP and RNA control the formation and dissolution of phase-separated Dhh1-rich droplets. **(A)** Dynamic light scattering and fluorescence microscopy show the formation of condensates upon addition of 5 mM ATP to a homogenous solution of Dhh1, most likely due to an increase of the intermolecular interactions upon ATP binding. **(B)** Upon ATP binding, the fraction of soluble Dhh1 monomer decreases to 33 ± 10 % with respect to a homogenous Dhh1 solution. **(C)** Dhh1 droplets formed in the presence of ATP and polyU can be dissolved by adding 36 µM RNase A, showing an additional level of regulation to form and dissolve the condensates. **(D)** Effect of ATP, AMP-PNP, ADP, AMP and GTP on the phase separation of Dhh1 quantified by evaluating the total droplet volume at each condition. **(E)** Fluorescence microscopy images of droplets in the presence of 2.5 mM ATP and 2.5 mM TNP-ATP, an ATP derivate which becomes fluorescent when bound to an ATP binding site. **(F, G)** Reversibility of droplet formation by removing ATP by fast-hydrolyzing apyrase. **(F)** Fluorescence microscopy images showing the kinetics of droplet dissolution upon addition of 1.2 µM apyrase. **(G)** The addition of 0 µM (green), 0.6 µM (blue), 1.2 µM (red) and 2.4 µM (black) of apyrase to droplets formed by 2 µM Dhh1 in presence of 5 mM ATP and 0.5 mg/ml polyU concentration-dependently dissolved the droplets.

The addition of 0.5 mg/ml polyU to a 2 µM Dhh1 (0.17 mg/ml) solution with 5 mM ATP / MgCl_2_ acted cooperatively with ATP binding in increasing protein-protein interactions and induced the formation of spherical, liquid-like protein condensates (Fig. 1, 2C). Assuming high and equal partitioning of the two components inside the droplets, the condensates contain, on average, 1 Dhh1 molecule per 750 nucleotides at these protein and polyU concentrations. The formed condensates could be dissolved by degradation of polyU upon RNase A addition (Fig. 2C), demonstrating one possible mechanism to regulate the disassembly of the droplets. Dhh1^48-425^ formed smaller droplets compared to full-length Dhh1 (Suppl. Fig. 3B), likely because the presence of the LCDs increases the intermolecular interactions between adjacent Dhh1 molecules. This is consistent with previous findings, that Dhh1^48-425^ forms fewer / no P-bodies in yeast cells (Hondele et al., 2019). Furthermore, only full-length Dhh1 was able to rescue P-body formation in a yeast strain deficient in two essential P-body components *(edc3△ lsm4△C)* whereas truncated Dhh1 lacking the LCDs did not (Protter et al., 2018).

We next investigated whether other nucleotides could promote phase separation of Dhh1 in the presence of polyU. To this aim, we analyzed a solution of 2 µM Dhh1 in presence of 0.1 mg/ml polyU and 5 mM of different nucleotides: ATP, its non-hydrolyzable analog adenylyl-imidophosphate (AMP-PNP), ADP, AMP or GTP (Fig. 2D, Suppl. Fig. 6). We observed formation of condensates only with ATP. This effect is consistent with the large intramolecular rearrangements that have been shown to occur in presence of ATP, but not with ADP and AMP-PNP (Cheng et al., 2005), and indicates that these conformational changes might be crucial for the protein to be able to undergo phase transition. Moreover, we further investigated the binding of ATP to Dhh1 inside the droplets by replacing 50 % of the ATP amount with 2,4,5-trinitrophenol adenosine triphosphate (TNP-ATP), whose fluorescence intensity increases upon interactions with the ATP-binding site (Stewart et al., 1998). Fluorescence emission was detected within the protein-rich condensates (Fig. 2E), indicating that ATP is bound to Dhh1 in the condensed phase.

These results demonstrate that the ATP-bound state of Dhh1 has a high propensity to undergo phase separation, suggesting that the removal of ATP, either by hydrolysis or by dissociation from the binding site, might promote the disassembly of the protein-rich condensates controlling their lifetimes. To directly test this, we introduced the enzyme apyrase, which hydrolyzes ATP with a higher reaction rate than Dhh1, into a solution of pre-formed droplets. Indeed, the addition of 0, 0.6, 1.2, 2.4 µM apyrase induced the concentration-dependent dissolution of the droplets within two hours (Fig. 2F and G).

Overall, our findings indicate that ATP binding increases the intermolecular protein-protein interactions of Dhh1. This mechanism is highly specific to ATP and the increase in protein-protein interactions acts synergistically with RNA binding in promoting the formation of condensates. The removal of either ATP or RNA is sufficient to dissolve the droplets providing distinct mechanisms to control the reversible assembly and disassembly of condensates.

### ATP hydrolysis, LCDs and RNA modulate the material properties and the maturation of the condensates

We next applied Differential Dynamic Microscopy (DDM) to investigate how droplet activity, LCDs and RNAs modulate the rheological properties and the maturation of the protein-rich condensates over time. DDM provides information on the dynamics of the system by analyzing a sequence of microscopy images taken in brightfield mode in a time interval of seconds to minutes (Fig. 3A). In analogy to dynamic light scattering, this technique reports on the sample dynamics by analyzing the fluctuations in the light scattered by the sample (see Materials and Methods) (Cerbino and Trappe, 2008; Giavazzi et al., 2009). Similar to particle-tracking strategies, the technique can be applied in combination with nano-sized particles of known size to quantify rheological properties. The size of the nanoparticles can be smaller than the diffraction limit as they do not have to be optically resolved in this technique. Here, we use nanotracers with a diameter of 25 nm (see Materials and Methods). When Brownian motion drives the dynamics of the tracers, the correlation function provides an effective diffusion coefficient *D*, from which the viscosity *η* of the medium can be estimated by the Stokes-Einstein relationship (*D = k*_*B*_*T/6πηR*).

**Figure 3.**
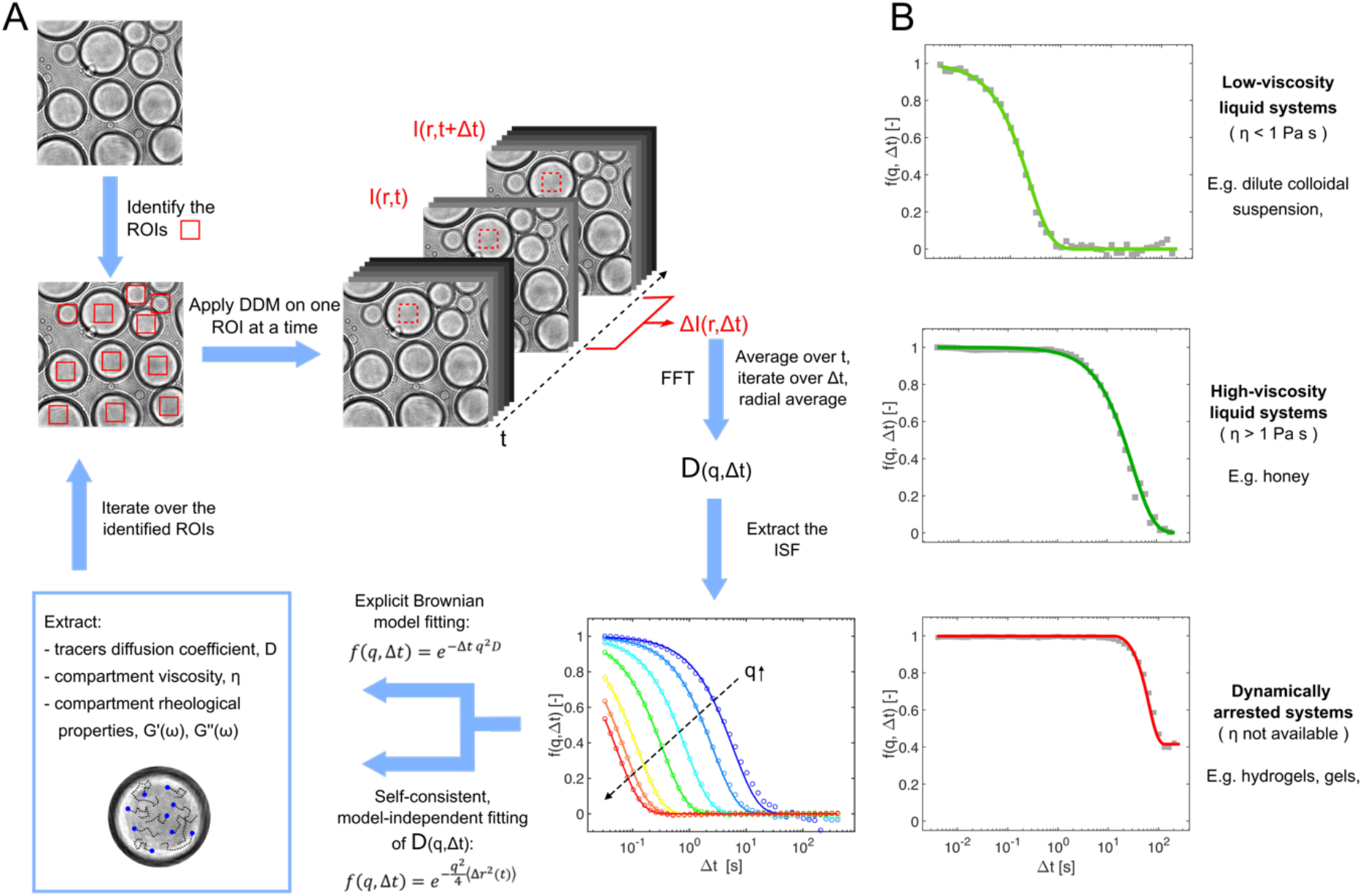
Differential Dynamic Microscopy to assess the material properties of biomolecular condensates. **(A)** Schematic representation of the DDM technique. **(B)** Representative intermediate scattering functions (ISF, f(q, △t)) for water, honey and a polymer-based hydrogel (from top to bottom).

Initially, we validated the method with mixtures of water and glycerol at different ratios and verified that the measured viscosities were consistent with values reported in the literature over a range spanning three orders of magnitude (0.001-1 Pa•s) (Suppl. Fig. 7A). Moreover, we demonstrated that DDM can be used to analyze highly viscous liquids such as honey (Fig. 3B) as well as dynamically arrested materials formed by liquid-to-gel transitions by monitoring the changes in the intermediate scattering functions (ISFs) during the gelation of the synthetic polymer polydimethylacrylamide (Fig. 3B, Suppl. Fig. 7B). In the case of dynamically arrested states, the viscosity cannot be computed using the Stokes-Einstein equation as the particle motion in such materials is not purely diffusive anymore. We further confirmed that the results in micro-sized gels generated *via* droplet microfluidics were consistent with bulk gels of the same material demonstrating the absence of confinement effects (Suppl. Fig. 7B) and the applicability of DDM to micron-size protein-/RNA-rich droplets.

After having identified polyU as an important factor to trigger the phase separation of Dhh1 (Fig. 2), we first applied DDM to investigate the effect of the Dhh1/polyU ratio on the droplet material properties (Fig. 4, Suppl. Fig. 8).

**Figure 4:**
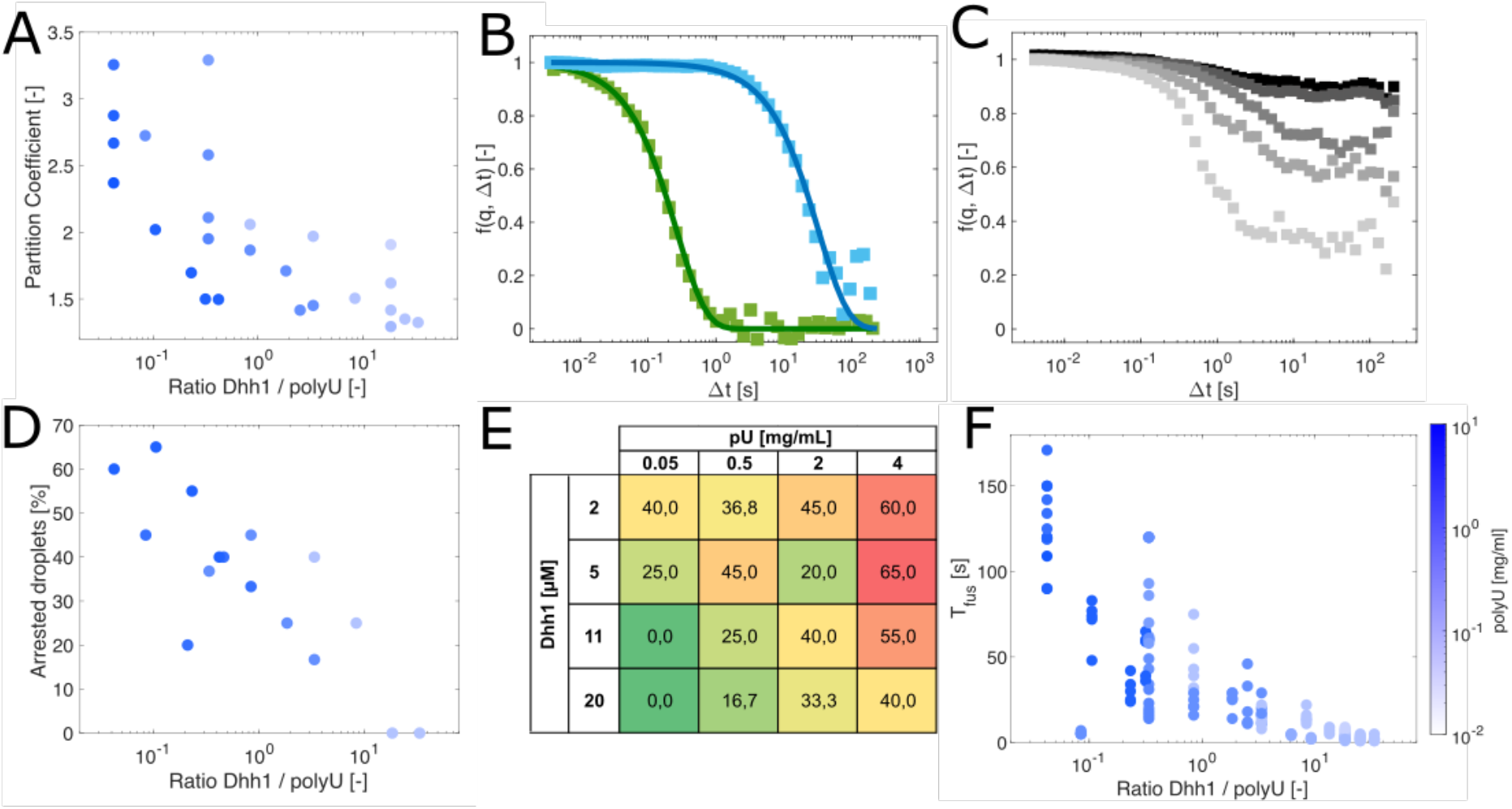
Effect of the Dhh1/polyU ratio on the droplet material properties. **(A)** A lower ratio of Dhh1 to polyU results in a higher partition coefficient (IntensityDhh1,droplets / IntensityDhh1,dilute) since the higher number of binding sites on polyU increases protein recruitment. **(B, C)** Representative ISFs f (q, △t) of liquid-like, low viscous (green curve) and highly viscous (blue curve) droplets (B), and dynamically arrested condensates formed in presence of various Dhh1/polyU ratios (C). **(D-F)** The increase of protein recruitment leads to a higher fraction of dynamically arrested droplets at low Dhh1/polyU ratios, as measured by DDM (D, E) as well as to an increase in the fusion time Tfus of the condensates (F). In panel (E), numbers indicate the fraction of arrested droplets in %, a high fraction of dynamically arrested droplets is highlighted in red, a low fraction is highlighted in green. The color bar in panel (F) describes the absolute polyU concentration used in (A, D, F).

We observed that the partition coefficient of Dhh1 measured by fluorescence intensity values outside and inside the condensates decreases with increasing Dhh1/polyU ratio (Fig. 4A), suggesting that high polyU concentrations offer a larger number of binding sites and therefore allow for a higher amount of protein to be recruited into the droplets. At high Dhh1/polyU ratios, the ISFs of condensates formed by full-length Dhh1 exhibited a single exponential decay characteristic of liquids (Fig. 4B), which is consistent with the results shown in Fig. 1. In contrast, at low Dhh1/polyU ratios, corresponding to high partition coefficients, condensates exhibited DDM correlation functions which were characterized by logarithmic decays and higher plateaus (Fig. 4C). These deviations are considered a hallmark of dynamically arrested systems such as gels and glass-like materials (Janssen, 2018). The fraction of these dynamically arrested droplets gradually increased when the Dhh1/polyU ratio was decreasing (Fig. 4D, E). This observation was confirmed by an increase in the droplet fusion time T_fus_ with a decrease in the Dhh1/polyU ratio (Fig. 4F).

These results show that the RNA molecules which are present in P-bodies are not only clients which are processed inside these compartments but contribute to the phase separation process as well as to the modulation of material properties of the condensates.

We next applied DDM to monitor the rheological properties of the condensates formed by full-length Dhh1, Dhh1^DQAD^ and Dhh1^48-425^ over five days of incubation (Fig. 5).

Based on the previous analysis of the material properties of the droplets at various Dhh1/polyU ratios, we selected a reference condition (11 µM Dhh1, 0.05 mg/ml polyU) to observe the maturation of the condensates over time starting from a population of 100 % low-viscous droplets (Fig. 5A, B). We estimated a viscosity in the range varying between 0.1 and 1 Pa•s, comparable to maple sirup, and only a modest change was observed during incubation (Fig. 5A). Similar viscosity values have been measured for condensates formed by proteins of the FUS family by active microrheology (Jawerth et al., 2019). Only on day 5 we observed the appearance of a second sub-class of condensates which exhibited arrested state (Fig. 5B). This drastic change in the rheological properties compromised the reversibility of the condensates upon dilution (Fig. 5C). These observations could be explained by the consumption of ATP over days. From previously determined hydrolysis rate (k_cat_ ≈ 0.001 1/s) (Dutta et al., 2011; Mugler et al., 2016) we estimated that the 5 mM ATP present in the mixture would be hydrolyzed after approximately 5 days, thereby interrupting the turn-over that likely keeps the droplets fluid. At this time point condensates could be stable even in the absence of ATP probably due to conformational rearrangements of Dhh1 and polyU occurred during incubation.

**Figure 5.**
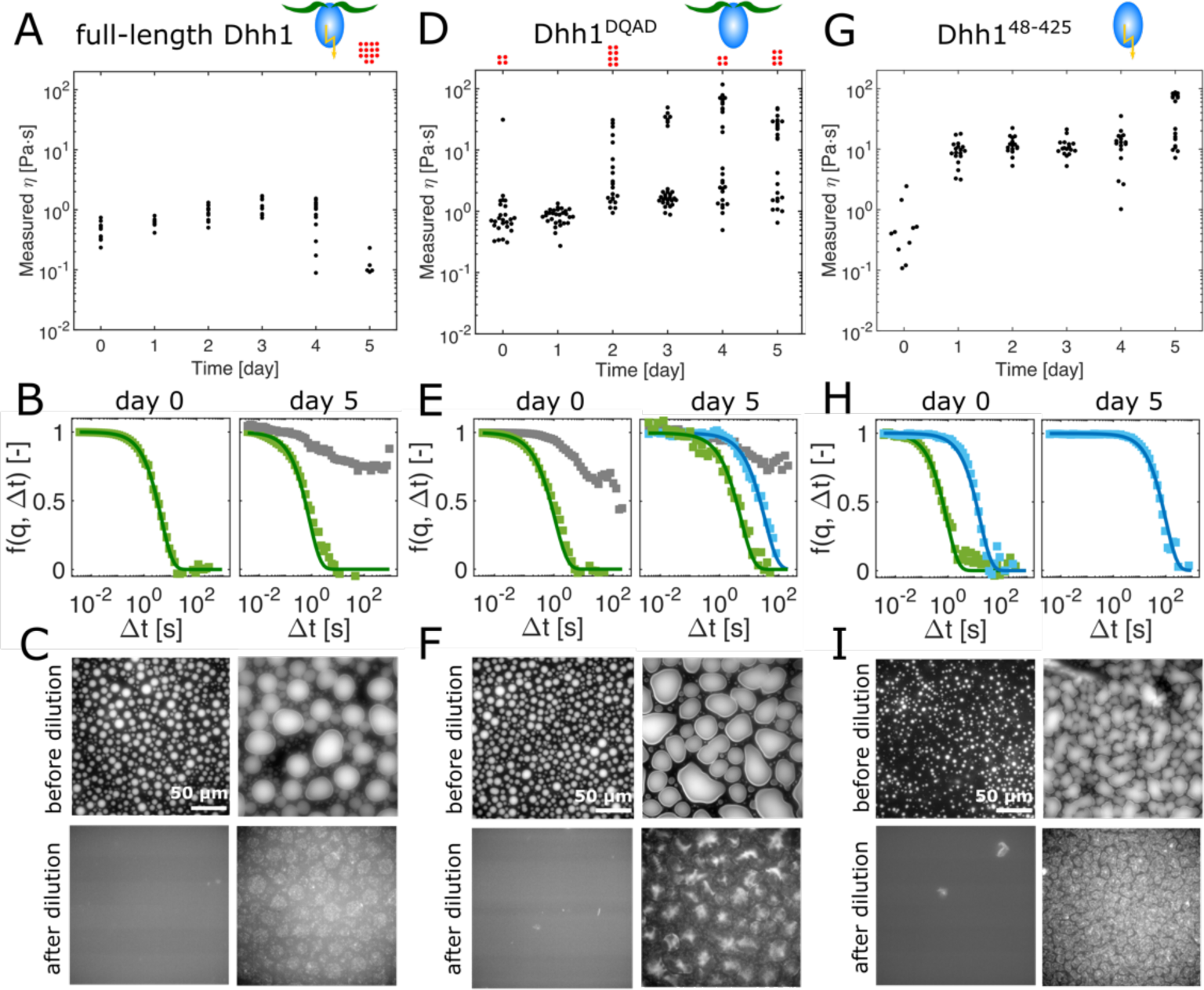
Material properties of full-length Dhh1, ATP-hydrolysis-deficient Dhh1^DQAD^ and the LCD-lacking variant Dhh1^48-425^. The viscosities of the condensates as well as the fraction of dynamically arrested gel-/glass-like droplets were measured over a time course of 5 days. Each dot represents a single droplet. **(A)** Full-length Dhh1 droplets remain liquid over several days (black dots) and only a fraction of the droplets becomes dynamically arrested on day 5 (red dots). For these droplets the viscosities could not be computed as they deviate from liquid-like behavior and are therefore displayed outside the graph. **(B)** Representative ISFs of droplets of full-length Dhh1 on day 0 (low-viscous, green) and day 5 (low-viscous, green; arrested, grey). **(C)** Droplets formed with full-length Dhh1 could be reversibly dissolved upon dilution on day 0 but not anymore on day 5, confirming the presence of non-liquid, or highly viscous structures. **(D)** In addition of low-viscous droplets, the catalytically deactivated Dhh1^DQAD^ variant showed a small fraction of dynamically arrested droplets already on day 0. The number of dynamically arrested droplets increased on day 2 and was accompanied by low- and high-viscous droplets. **(E)** Typical ISFs of Dhh1^DQAD^ droplets, showing low-viscous (green) and arrested droplets (grey) on day 0 and low-, high-viscous (blue) and arrested droplets on day 5. **(F)** Also the Dhh1^DQAD^ droplets could not be dissolved by dilution on day 5, in contrast to full dissolution on day 0. **(G)** Droplets formed in the presence of Dhh1^48-425^ showed an increase of viscosities of about one order of magnitude with respect to Dhh1 after day 1. No dynamically arrested droplets were observed over the time course of 5 days. **(H)** Representative ISFs of Dhh1^48- 425^ droplets with low viscosity on day 0 (green) and of droplets with high viscosity observed after day 1 (blue). **(I)** Also Dhh1^48-425^ droplets could be dissolved on day 0 but not on day 5

It has previously been proposed that the removal of enzymatic activity could lead to hardening and irreversibility of biomolecular condensates. For instance, ATP-hydrolysis-deficient variants of Dhh1 (Carroll et al., 2011; Mugler et al., 2016), Ded1 (Hilliker et al., 2011) or DDX3X (Valentin-Vega et al., 2016), and Vasa (Xiol et al., 2014) form constitutive granules inside cells, even in the absence of stress. In contrast, ATP hydrolysis keeps these granules dynamic (Carroll et al., 2011; Hondele et al., 2019).

Consistent with this hypothesis, we observed that the catalytically inactive Dhh1^DQAD^ mutant exhibited a significant fraction of condensates with ISFs corresponding to an arrested state shortly after formation (Fig. 5D, E). Moreover, already on day 2 of incubation we observed several subclasses of condensates characterized by different viscosity values. For this variant, the change in rheological properties also compromised the reversibility of the condensates (Fig. 5F). We note that Dhh1 and Dhh1^DQAD^ have very similar behaviors with respect to phase separation (Suppl. Fig. 2, 4), but drastically differ in terms of their material properties.

To rule out a potential effect of the tracers, we performed experiments also in the absence of nanoparticles, analyzing the scattering signal that directly originates from the macromolecules. Although in this case no viscosity value can be derived, as the size of the scattering species is not known, the shape of the autocorrelation function was consistent with and without nanoparticles, demonstrating the same qualitative behavior and the transition from liquid to an arrested state (Suppl. Fig. 9A).

We next analyzed the role of the LCDs on the maturation of the droplets by investigating the behavior of the truncated variant Dhh1^48-425^ (Fig. 5G). Although these droplets showed similar phase separation behavior as full-length Dhh1 (Suppl. Fig. 3), the condensates exhibited higher values of viscosities already after one day of incubation, and the viscosity increased by one order of magnitude over time. On day 5 we observed the presence of two sub-classes of condensates, one of which exhibited remarkably high viscosity values in the order of 80-100 Pa•s. Such high viscosity values are consistent with those of condensates formed by Laf-1, Whi3 or GAR-1△N, as measured by passive microrheology (Taylor et al., 2016). However, no condensates with ISFs characteristic of arrested materials were observed (Fig. 5H). Also in this case, the reversibility of the condensates upon dilution was significantly impaired on day 5 (Fig. 5I). These results show the important role of LCDs not only in modulating phase transition but also in maintaining fluid-like properties over time. This result is not intuitive since the promotion of phase separation requires attractive interactions, which however often results in high viscosity values. By contrast, the truncated variant Dhh1^48-425^ shows phase-separated droplets smaller in size (Suppl. Fig. 3B) but exhibiting higher viscosity values.

### Structured RNA induces dynamically arrested states which can be partially rescued by stimulating the turnover of the droplet material

So far, our analysis involved the RNA mimic polyU, which contains only one type of nucleotide leading to linear RNA molecules. To mimic better physiologically occurring RNAs we substituted the polyU with an *in vitro*-transcribed, structured, 600 nt RNA (Suppl. Fig. 10A and Materials and Methods).

Upon addition of this structured RNA, Dhh1 again formed condensates instantaneously after mixing, although a higher protein concentration was required (11 µM instead of 2 µM Dhh1 in the presence of polyU) (Suppl. Fig. 10B). Analysis of the material properties by DDM revealed that these droplets contained a large subpopulation of dynamically arrested droplets (low-viscous, 48±4%; high-viscous, 4 ± 2 %; and dynamically arrested, 47 ± 4%) compared to the system containing polyU (low-viscous, 100 %) (Fig. 6A, 5B, 7B), even directly after formation. This loss of droplet fluidity in presence of structured RNA was confirmed by FRAP experiments, which showed that the fraction of mobile molecules (defined as the percentage of recovery after 30 s) was about 88 ± 2 % directly after formation (Fig. 6B) and decreased rapidly to around 5 ± 5 % after 2 h. Droplets formed at the same concentration of polyU showed higher recovery after the same time (Suppl. Fig. 10C, Fig. 7C). Furthermore, in the presence of structured RNA, the condensates underwent a dehydration process over time that led to shrinking and changes in morphology from a spherical to an irregular shape (Fig. 6B, 7D, E). In addition, these condensates could not be dissolved by dilution, confirming their non-liquid nature (Fig. 6C).

**Figure 6.**
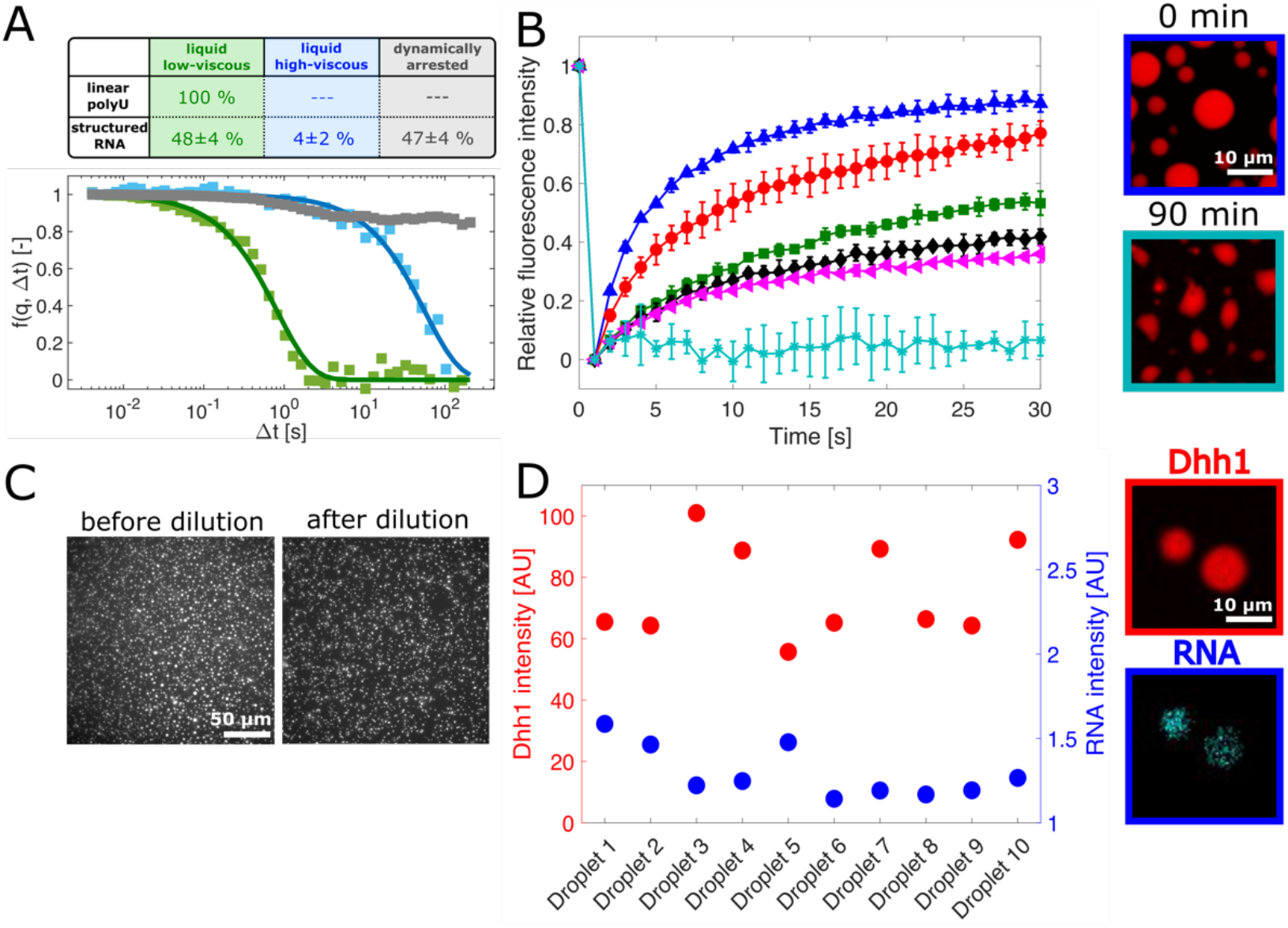
Effect of structured RNA on the dynamics of the condensates. **(A)** The addition of 0.05 mg/ml *in vitro* transcribed, structured RNA (600 nucleotides) to a solution of 11 µM wild-type Dhh1 and 5 mM ATP / MgCl2 lead to the formation of droplets with heterogeneous material properties, as revealed by DDM measurements, exhibiting a high fraction (47±4%) of glass-/gel-like droplets, in contrast with droplets formed in presence of the same amount of polyU (0 %). Intermediate scattering functions of exemplary droplets (low-viscous, green; high-viscous, blue; dynamically arrested, grey). **(B)** Fluorescence recovery after photobleaching (FRAP) measurements showed a decrease in recovery over a time course of 2 h (blue - 15 min, red - 30 min, green - 45 min, black - 60 min, magenta - 90 min, light blue - 120 min). Simultaneously, the morphology of the droplets changed from an initially spherical to an irregular shape. **(C)** The condensates formed in presence of structural RNA cannot be dissolved by dilution. **(D)** Different droplets imaged at the same time point exhibited different ratios of mCherry-tagged Dhh1 and Fluorescein-12-labeled RNA intensities, corresponding to fluctuations of the protein / RNA concentrations which might explain the presence of droplet subpopulations with different material properties.

**Figure 7.**
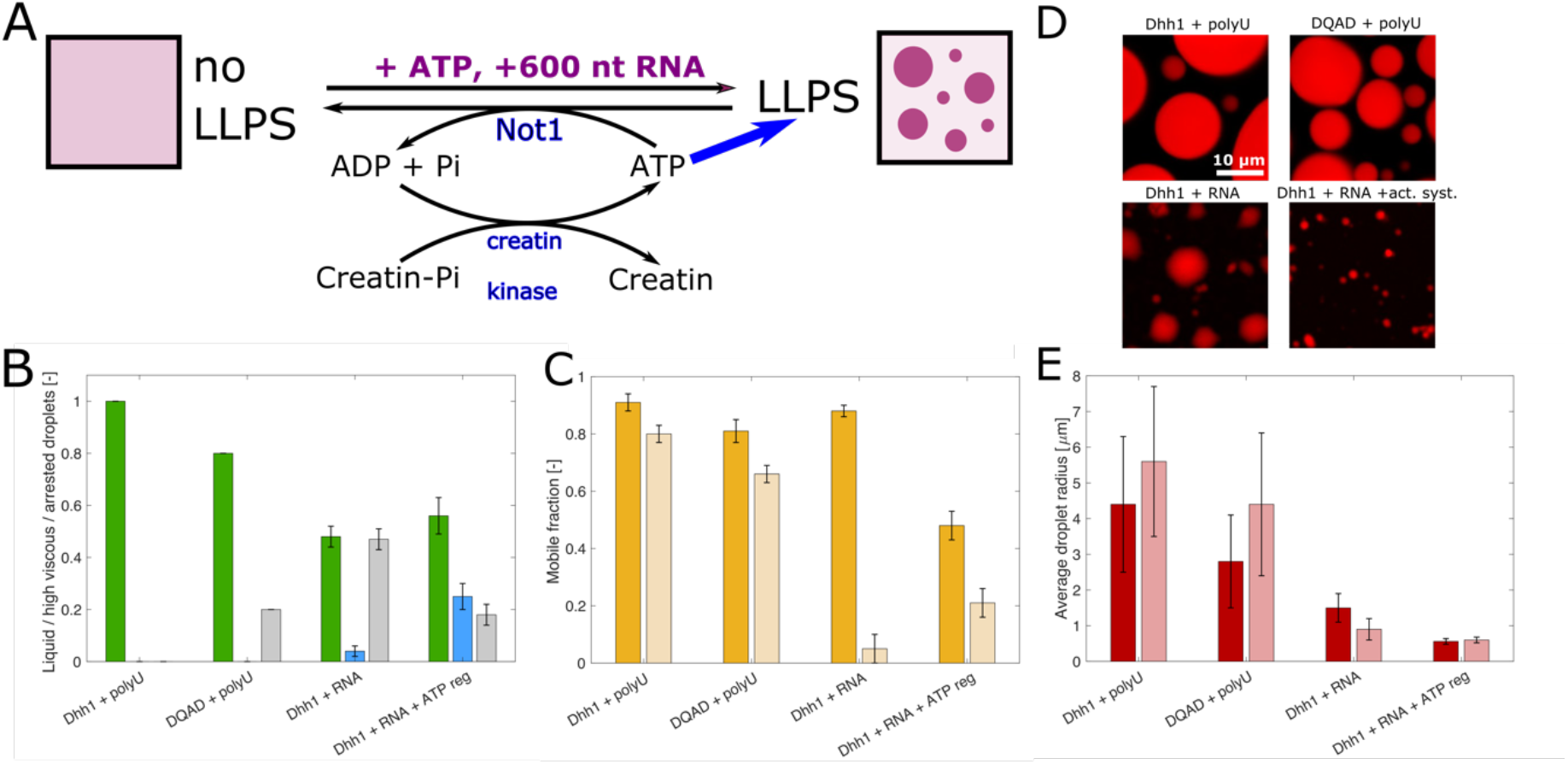
Effect of droplet turnover on material properties of biomolecular condensates. **(A)** Schematic illustration of the ATP-hydrolysis-regeneration system: addition of ATP and RNA to a homogenous Dhh1 solution induces droplets formation, while introduction of Not1 stimulates the ATPase activity of Dhh1 and promotes droplets dissolution. ATP is restored by creatin kinase, which phosphorylates ADP using creatin phosphate as phosphate donor. **(B)** Fractions of the different droplet sub-populations characterized by DDM: low-viscous liquid (green), high-viscous liquid (blue) and dynamically arrested (grey). **(C)** Mobile fraction extracted from FRAP measurements at time 0 (dark yellow) and after 90 min incubation (light yellow). In presence of polyU, the mobile fraction was about 91 ± 3 % and remains almost constant over time. A similar behavior was observed for the Dhh1^DQAD^ variant. When polyU was replaced with structured RNA, the mobile fraction decreased to 5 ± 5 % over time, and this decrease could be partially rescued when coupled to an active system. **(D)** Confocal images of condensates after 90 min incubation. **(E)** Average size of the condensates at time 0 (dark red) and after 90 min incubation (light red). Over time, the condensates formed in presence of full-length Dhh1 and Dhh1^DQAD^ increased their size due to droplet coalescence. By contrast, condensates containing structured RNA decreased in size due to a dehydration process, which could be inhibited by increasing droplet activity.

**Figure 8:**
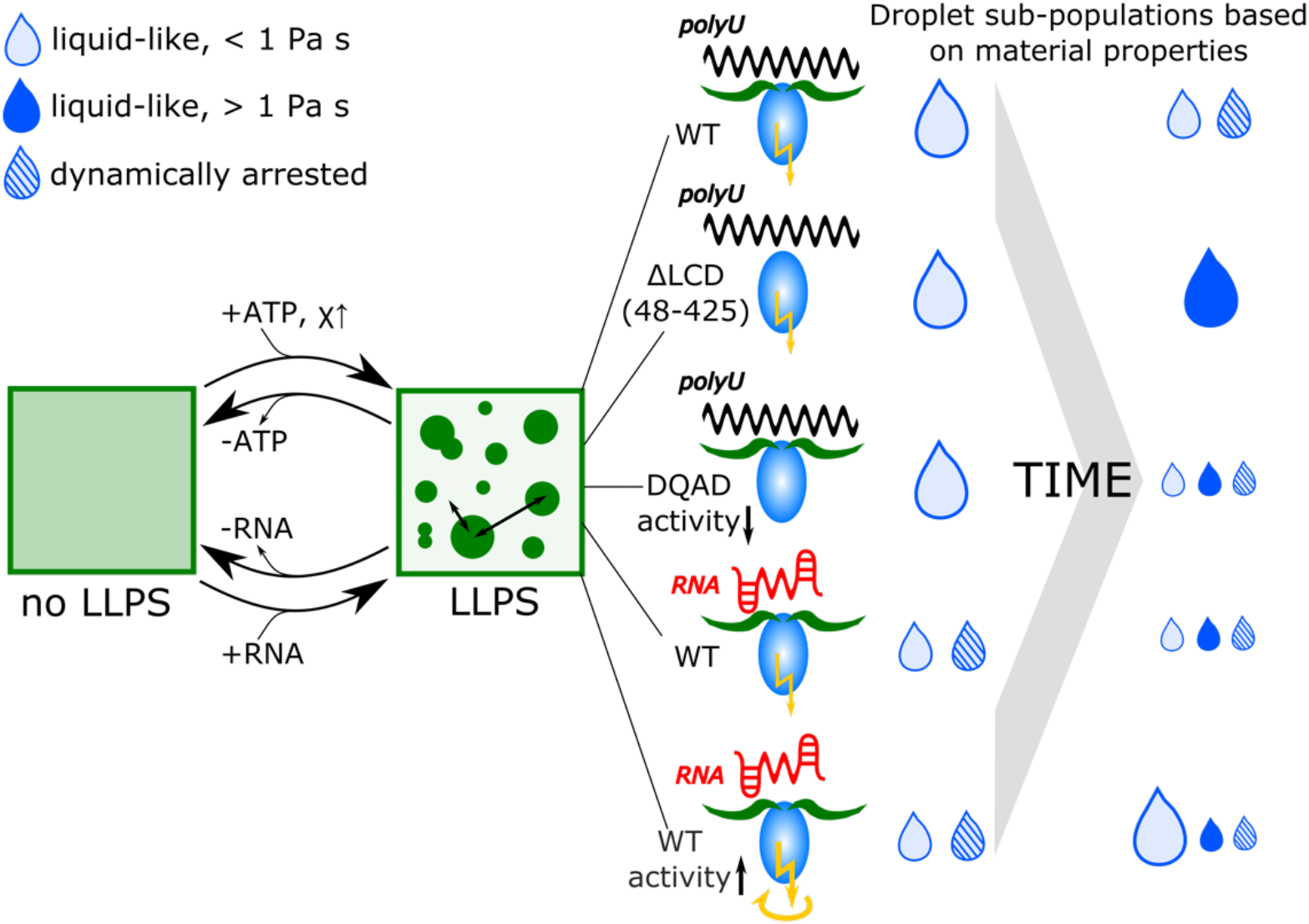
Summary of the dynamics of phase transition and the effect of various modulators. In absence of a stimulus, Dhh1 remains soluble even at high concentrations. Upon addition of ATP and RNA, dynamic liquid-like droplets form, which exchange molecules with their surroundings. The addition of only ATP is sufficient to increase the intermolecular interactions to form aggregate-like condensates. This phase transition is reversible by removing either ATP or RNA from the system. Droplets formed by full-length Dhh1 remain liquid-like over several days, while the deletion of the LCDs (Dhh1^48-425^) as well as the depletion of the active domain of Dhh1 (Dhh1^DQAD^) induce a decrease of the dynamicity over days (light blue drop – liquid, low viscous; dark blue drop – liquid, high viscous; striped, blue drop, dynamically arrested). Molecular features embedded in the protein itself represent therefore a first important level of droplet dynamicity. In addition, external stimuli responsible for triggering phase separation also modulate the material properties over time. Exchanging the unstructured polyU with structured RNA immediately arrests the droplet dynamics. This arrest can be inhibited by stimulating the turnover of material between the condense and the dilute phase by promoting ATPase activity (coupled with a ATP-regenerating system).

The heterogeneity of droplet material properties could occur due to a non-homogeneous distribution of Dhh1 and RNA molecules among the condensates. Consistent with this hypothesis, measurements of the fluorescence intensities of mCherry-tagged Dhh1 and Fluorescein-12-labeled RNA within ten different condensates (Fig. 6D, Suppl. Fig. 10D) showed differences of almost 40 % in the protein signal and of 30 % in the RNA signal. This heterogeneity in composition indicates low dynamics of material exchange between the different condensates after their formation, suggesting that stronger protein-protein, RNA-RNA, and protein-RNA interactions lead to the formation of heterogeneous networked fluids.

We next attempted to restore droplet dynamicity also in presence of structured RNAs by promoting the turnover of the droplet-forming / droplet-dissolving material.

To this aim, we stimulated the ATPase activity of Dhh1 by adding Not1 into the system, which induces droplet dissolution. Simultaneously, we induced droplet re-assembly by converting the resulting ADP into ATP by addition of creatine kinase (using creatine phosphate as phosphate donor). Using this active system, the reversible formation and dissolution of droplets was continuously promoted by a rapid exchange of proteins and RNA molecules between the condensed and diluted phase, until all creatine phosphate (our “fuel”) was consumed (Fig. 7A). Under these conditions, the arrested droplets could partially be rescued and the average number of droplets with arrested dynamics was significantly decreased to about 18 ± 4 %, which corresponds to a reduction of 30 % compared to the non-active system (Fig. 7B).

These findings were consistent with the higher mobile fraction compared to the non-active system measured by FRAP after 2h (Fig. 7C).

The increase of enzymatic activity of the droplets formed in presence of the ATP-hydrolysis-regeneration system did not only affect the material properties but also the size of the droplets (Fig. 7D). In general, the droplets formed in presence of full-length Dhh1 and polyU were the largest, while the ones in presence of full-length Dhh1, structured RNA and the ATP-hydrolysis-regeneration system were the smallest (Fig. 7D). While the droplets formed in presence of polyU undergo coalescence within the first 90 min after formation, the droplets formed in presence of structured RNA decreased in size, most likely due to water loss while approaching equilibrium. This decrease of size could be rescued by introducing the ATP-hydrolysis-regeneration system where it remained constant over the first 90 min after formation (Fig. 7E).

These results show that biomolecular condensates formed via active phase-separated systems which are constantly turned over by biochemical reactions remain fluid over longer time scales, indicating that the coupling of biochemical reactions with phase separation provides a mechanism to prevent or at least delay droplet maturation.

## Discussion

Here, we have investigated mechanisms controlling the material properties of biomolecular condensates consisting of the DEAD-box ATPase Dhh1, a simplified *in vitro* model of P-bodies. To this aim, we introduced Differential Dynamic Microscopy (DDM), which allowed us to probe the dynamics of the condensates and distinguish between liquid-like and arrested states. The DDM analysis can be performed *in situ*, even in the absence of tracer particles and at the single condensate level. The latter aspect was very important in revealing that Dhh1 condensates exhibit a distribution of material properties that can be modulated by various factors.

To investigate the molecular basis underlying this rich behavior, we first characterized the effect of multiple factors (LCDs, ATP and RNA) on the reversible formation and dissolution of the droplets (Fig. 1 and 2). We observed a hierarchy of intermolecular interactions (Schmit et al., 2020) encoded by these different modulators. Based on the emerging “stickers and spacers” model (Choi et al., 2020), the molecular architecture of the interacting biomolecules can be divided into stickers that mediate the intermolecular interactions driving LLPS and spacers which modulate other chain properties. In the absence of RNA, we propose that the LCDs of Dhh1 act as stickers mediating weak multivalent interactions, while the globular RecA domains represent spacers. Indeed, the C-terminal LCD of Dhh1 undergoes liquid-liquid phase separation on its own (data not shown). In the absence of RNA, we observed phase separation only at low salt concentrations, which strengthen attractive LCD-LCD interactions. However, in presence of RNA, the RNA-protein interactions dominate over the weak multivalent interactions of the LCDs and therefore these heterotypic interactions become the sticky element which determines the phase behavior. These stronger RNA-protein interactions are less sensitive to changes in the salt concentration (Sup. Fig. 2).

Surprisingly, the DDM analysis showed that in case of Dhh1, the LCDs do not only promote LLPS, but they are also important in maintaining droplet fluidity over time. Indeed, droplets formed by the LCD-lacking variant Dhh1^48-425^ exhibited higher viscosities than droplets formed in presence of full-length Dhh1. This change in viscosity could be due to the increased number of accessible binding sites on the RNA molecules for the smaller variant protein compared to the full-length Dhh1, which in turns lead to the recruitment of larger amounts of protein within the condensates (Fig. 5A, G). This suggests that the LCDs may switch from being a driver of phase separation in absence of polyU to being a modulator of the material properties in presence of polyU. We propose that this is an additional important function of LCDs in biomolecular condensates.

Furthermore, we identified the structure of RNA as another important external modulator of the material properties of the condensates. Specifically, structured RNA drastically decreased the dynamics of a large sub-population of the Dhh1 condensates. The presence of structured RNA in these condensates increases the complexity of the system. RNA base pairing leads not only to a complex three-dimensional structure of the RNA itself, but allows also for intermolecular RNA-RNA interactions, which are further promoted by the high RNA concentration inside the droplets. While some of these interactions could be transient and dynamic, others could be energetically favored and stabilized over time, and might thereby contribute to the formation of networked droplets and droplet hardening.

Importantly, we find that the richer ensemble of intermolecular protein-protein, protein-RNA and RNA-RNA interactions leads to the presence of droplet subpopulations characterized by different material properties. This highlights the importance of single-droplet techniques to analyze these heterogeneous populations. We have demonstrated that the broad distribution of material properties correlates with differences in the protein / RNA concentration inside the droplets (Fig. 5), which should therefore be seen as networked condensates characterized by slow exchange of material between the condensed and the diluted phase.

Moreover, our results reveal that the mobility of different molecular species within the condensates can vary significantly, and these differences can be probed by complementary techniques. For instance, DDM shows the presence of a large fraction of arrested droplets already shortly after droplet formation (Fig. 6A). By contrast, FRAP analysis on the same samples indicates high diffusivity of the proteins (Fig. 6B). The three-dimensional network likely consists of structured RNA molecules that recruit Dhh1 molecules. This network contains gaps which can be initially filled by highly diffusive Dhh1 molecules. The release of these molecules over time leads to the shrinkage of the droplets which are now formed by the network of RNA molecules containing only the Dhh1 molecules strongly attached to them, as indicated by the low FRAP recovery and the droplet irreversibility.

We have further demonstrated that droplet activity is an important mechanism to modulate their material properties. In this work, we refer to “active droplets” as the formation and degradation of our condensates depends on biochemical reactions that simultaneously generate components characterized by low and high propensities to phase separate (Dai et al., 2020; Tena-Solsona et al., 2018; Weber et al., 2019; Zwicker et al., 2017) (Fig. 7A, 8).The Dhh1 system allows us to investigate the effect of activity on the droplet material properties on several levels, since the intrinsic enzymatic ATPase activity of Dhh1 can be modulated in different ways.

Condensates formed in presence of full-length Dhh1 and polyU exhibit high fluidity, despite the low intrinsic propensity to hydrolyze ATP (Mugler et al., 2016). We note that under these conditions the hydrolyzed ATP is not actively regenerated, and our results suggest that ATP consumption is therefore the limiting factor determining the dynamic arrest of the droplets on day 5.

Condensate fluidity can be decreased when ATP hydrolysis is inhibited by exchanging a single glutamate (E) to glutamine (Q) in the DEAD box of the protein (Dhh1^DQAD^) (Carroll et al., 2011; Mugler et al., 2016). Droplets formed in presence of this variant rapidly form large populations of highly viscous and gel-/glass-like droplets showing the importance of enzymatic activity in their interior.

While condensates formed in presence of full-length Dhh1 and polyU exhibit highly liquid properties, condensates formed in presence of structured RNA are largely dynamically arrested, even shortly after their formation. Increasing the Dhh1 activity employing an ATP hydrolysis-regeneration system (Fig. 7A) promotes liquidity and partially rescues the dynamic arrest of the droplets (Fig. 7B). This is likely due to the accelerated turnover of the droplet material between the dispersed and the diluted phase, which keeps the system out of equilibrium. However, this increase of droplet turnover was not sufficient to fully liquefy all droplets, indicating the importance of structured RNA in tuning the material properties of condensates and in being a critical driver for their dynamic arrest (Fig. 7B).

Since processing bodies are part of the cellular metabolism and are therefore intrinsically out of equilibrium systems, this role of biochemical reactions in keeping fluid-like properties can be important also in the cellular context to maintain functional P-bodies and counteract the effect of client mRNAs, which may otherwise compromise their dynamics.

Furthermore, our study shows that liquid-like condensates can “age” towards dynamically arrested materials over time, consistent with recent findings demonstrating that the relaxation time of some condensates increases with age, in analogy with a glass-forming system (Jawerth et al., 2019). For all the investigated Dhh1 variants, the condensates exhibited either an increase in the viscosity over time or a transition towards a gel / glass state. Interestingly, we did not observe formation of protein aggregates or amyloids, in contrast with other phase-separated condensates formed in presence of FUS or hnRNPA1 (Molliex et al., 2015; Patel et al., 2015).

Since this ageing occurs without any change in extrinsic variables such as temperature, the dynamic arrest is likely governed by rearrangements of LLPS-driving molecules inside the droplets, which in turn modulate intermolecular interactions, therefore driving the relaxation of the initial kinetically trapped droplets into an arrested state.

These results show that biomolecular condensates are carefully regulated on several levels by nature not only to reversibly assemble and disassemble in the presence of suitable triggers but also to maintain the appropriate level of fluidity required for their function. Some of the regulating features are embedded in the architecture of the scaffold protein itself (e.g., LCDs), while other factors are extrinsic (e.g., RNA molecules with different structures).

## Materials and Methods

### Protein expression and purification

Expression and purification of mCherry-tagged Dhh1 (mCh-Dhh1), Dhh1^48-425^, Dhh1^DQAD^ and non-tagged MIF4G-Not1 was performed as previously described (Mugler et al., 2016). Briefly, competent *Escherichia coli* BL21-Gold (DE3) strains were transformed via heat shock at 42°C with plasmids carrying the genes for Dhh1 (pKW3631), Dhh1^48-425^ (pKW4063), Dhh1^DQAD^ (pKW3632) or Not1^MIF4G^ (pKW3469). Each plasmid was carrying sequences for a 6x His tag and ampicillin resistance. Cells were cultured in LB medium at 37 °C and protein expression was induced by addition of 0.5 mM (0.2 mM for Not1^MIF4G^) isopropyl-beta-D-1-thiogalactopyranoside (IPTG). After harvesting, cells were resuspended in lysis buffer (pH 7.5, 300 mM NaCl, 50 mM Tris, 10 mM imidazole, 10 % glycerol) and lysed by sonication. Protein purification was performed via affinity chromatography using Ni^2+^ charged Fast Flow Chelating Sepharose (GE Healthcare) as stationary phase. This step was followed by size exclusion chromatography on a Superdex 75 column (GE Healthcare) using a solution at pH 7.5, 300 mM NaCl, 25 mM Tris, 2 mM 2-Mercaptoethanol and 10 % glycerol as elution buffer. Purified fractions were pooled, concentrated and flash-frozen in liquid nitrogen. The phase diagram of Dhh1 was typically analyzed in buffer b-150, a 30 mM HEPES-KOH buffer at pH 7.4 supplied with 150 mM KCl and 2 mM MgCl_2_.

### *In vitro* transcription and labeling of RNA

For *in vitro* transcription, a construct was designed consisting of a 6 x 100 nucleotide repeat of actin mRNA interspaced by 6 different restriction site linkers (Suppl. Fig. 10) and commercially synthesized (GeneWiz). After amplification in *E. coli* bacteria and isolation from the cells using a QIAprep Spin Miniprep Kit (Qiagen), the plasmid was linearized by restriction enzyme digestion at the last restriction site RS6, by adding 1 µl (20 units) of BamHI restriction enzyme (New England Biolabs) to 1 µg DNA. The linearized DNA was purified on a 1% agarose gel, isolated from the gel and *in vitro* transcribed using a MEGAshortscript™ Transcription Kit (ThermoFisher Scientific) according to the manufacturer’s instructions. For labeling, 0.9 mM of fluorescein-12-labeled UTP (Jena Bioscience, Germany) was added. The mixture was incubated overnight at 37 °C and the resulting RNA was purified by ethanol precipitation.

### Microscopy

For microscopy analysis 20 µl of samples were transferred to a 384-well plate (Brooks, Matriplate) and imaged by either wide-field microscopy or confocal fluorescence microscopy. Analysis by wide-field microscopy was performed on an inverted epi-fluorescence microscope (Nikon Eclipse Ti-E) equipped with a 60x NA 1.4 oil objective (Nikon), an LED light source (Omicron LedHUB Light engine) and an Andor Zyla sCMOS camera. Confocal microscopy images were acquired with an inverted epi-fluorescence microscope (Leica TCS SP8) equipped with an 63x NA 1.4 oil objective (Leica), a Laser unit for confocal acquisition (AOBS system) and a sCMOS camera (Hamamatsu Orca Flash 4.0).

The size distributions of the droplets were reconstructed from the images acquired by optical and fluorescence microscopy via an in-house program written in Matlab (Suppl. Fig. 3B).

### Size exclusion chromatography to determine the soluble monomer concentration

The soluble Dhh1 concentration (C_S_) was measured by UV absorbance after removing the protein-rich droplets by centrifugation (10 min at maximum speed) on a bench top centrifuge and running the supernatant on a Superdex 200 size exclusion column (GE Healthcare).

### Dynamic Light Scattering

We used dynamic light scattering to measure the hydrodynamic radius of the protein and the resulting condensates. 100 µl of sample were prepared containing 5 µM Dhh1 in absence or presence of 5 mM ATP and 25 mM GTP. The sample was measured on a Zetasizer Nano ZS instrument (Malvern) in 173° backscattering mode in a quartz cuvette (Hellma Analytics, Germany).

### Fluorescence Recovery After Photobleaching (FRAP)

FRAP experiments were performed on the confocal microscope described above. Droplets were bleached by focusing a 561 nm laser light on a circular area with a diameter equal to about one tenth of the total droplet diameter. Image analysis, including background subtraction, correction of bleaching during recovery and normalization to pre- and postbleach intensity was performed via an in-house program written in Matlab.

### Differential Dynamic Microscopy (DDM)

A sample volume of 20 µL was introduced in at least three wells of a 384-well plate with quartz bottom (Matriplate, Brooks Life Sciences, USA). 4 µL of fluorescently labelled nanotracers with a diameter of 25 nm (29-00-251 micromer®-greenF, Micromod Partikeltechnologie GmbH, Rostock, Germany) were added to each well. The samples were stored at 4°C over several days and equilibrated to room temperature before the measurement.

Stacks of bright field images were acquired on a Ti2-U epi-fluorescence inverted microscope (Nikon) equipped with an sCMOS camera (Zyla 4.2P-CL10, Andor, UK) and with a 60x magnification water objective (CFI Apochromat NIR 60X W, Nikon, Japan, NA = 1.0). Sequences of N = 1000-4000 images of 512 × 512 pixels (corresponding to 55.3 × 55.3 µm^2^) were acquired both at high frequency (250 frames per second) and low frequency (4 fps) to capture the short- and long-time sample dynamics, respectively. The exposure time was kept fixed throughout the experiments at 1 ms. The images were processed and analyzed with a custom written Matlab code. The size of the selected regions of interest varied between 64 × 64 and 256 × 256 pixels, depending on the samples.

We verified that the DDM measurements were not affected by confinement effects, by wetting or by the size of the nanoparticles, by performing a large number of control experiments with condensates of different sizes (Suppl. Fig. 7B, D) and with tracers of different sizes (Suppl. Fig. 9).

Furthermore, we verified the absence of confinement effects that may arise due to the micron-sized range of the investigated phase-separated protein compartments by comparing a model polydimethylacrylamide (PDMA) hydrogel matrix in bulk (in a 384-well plate) and within compartments generated by droplets microfluidics. Immediately after their generation in the microfluidic chip, the gelling droplets were collected in a glass capillary. Prior to DDM imaging, the capillary was sealed with epoxy glue to prevent evaporation and the accidental motion of the droplets (Suppl. Fig. 7B).

## Acknowledgements

The authors gratefully acknowledge financial support from the Swiss National Science Foundation (grants 205321_179055 (PA), 31003A_179275 (KW) and CRSII5_193740 (KW)), the Synapsis Foundation (PA) and the Claude and Giuliana Foundation (PA). We are grateful for the help of Cora De Gol, Dany Liu, Marie Kopp, Andreas Küffner, Dr. Lenka Faltova and Dr. Umberto Capasso Palmiero with experiments, and for the assistance and support of Dr. Joachim Hehl and Dr. Justine Kusch (ScopeM, ETH Zurich) for confocal microscopy and FRAP experiments. We gratefully acknowledge Giovanni Savorana (Dep. Civil, Environmental and Geomatic Engineering, ETH Zurich) and Paolo Edera (Department of Physics, University of Milan) for helpful discussions, as well as Prof. Eric Dufresne (Dep. of Materials, ETH Zurich) for critical reading of the manuscript.

