## Supplementary Information for "Dynamic arrest and aging of biomolecular condensates are regulated by low-complexity domains, RNA and biochemical activity"

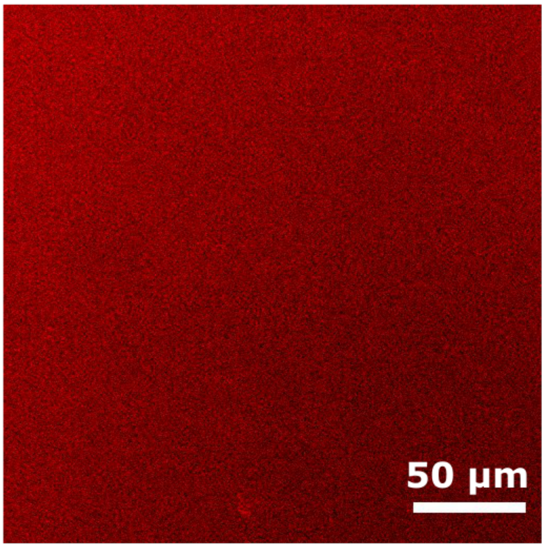


**Figure S1:** 500 μM Dhh1 solution does not undergo phase separation in absence of ATP and RNA, showing the requirement of both modulators to trigger droplet formation.

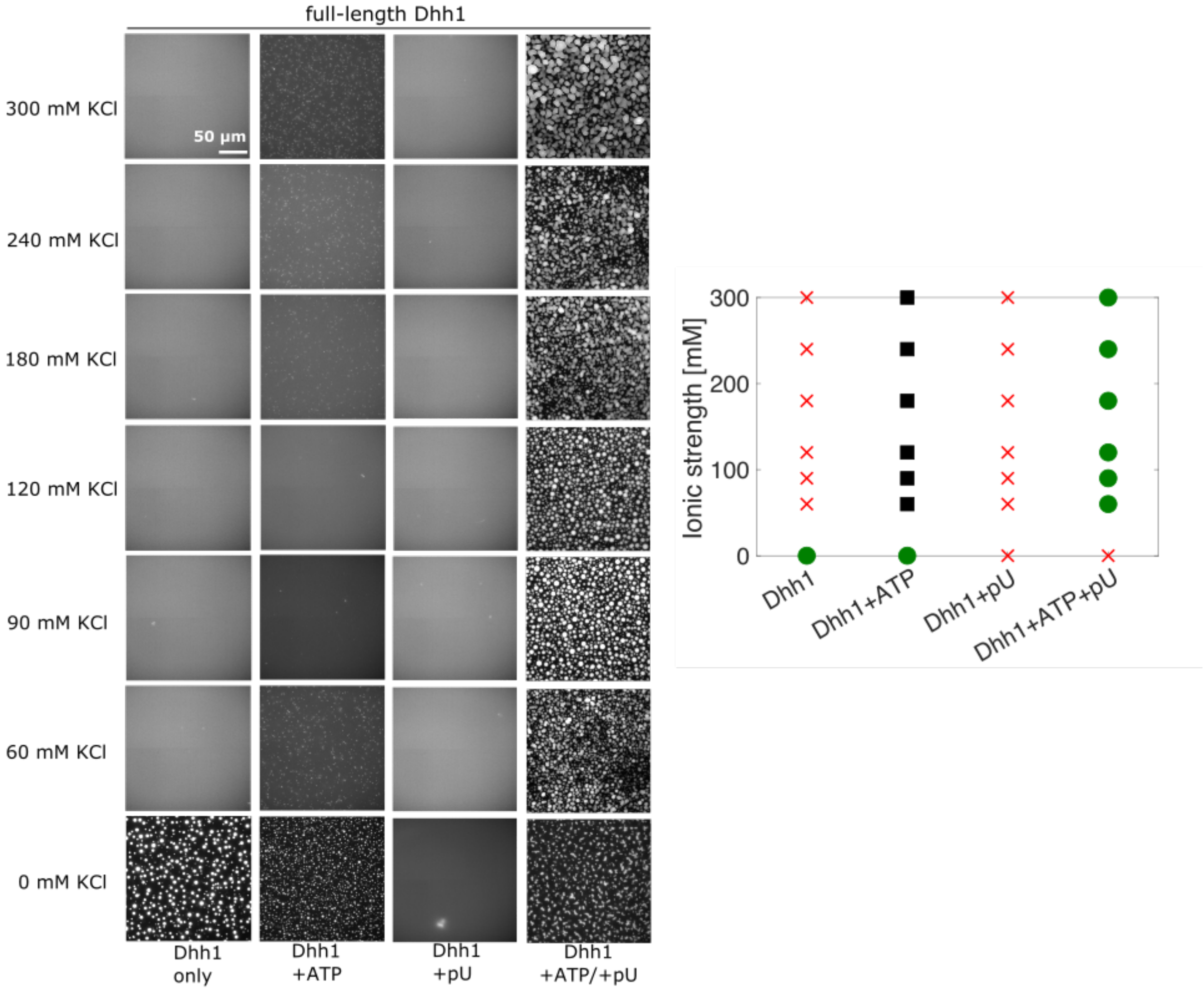


**Figure S2:** Phase diagram of full-length Dhh1 in absence / presence of ATP, polyU and ATP/polyU at increasing KCl concentration. Left: fluorescence microscopy images; right: phase diagram. Red crosses – no phase separation, green circles – phase separation, black squares – small aggregates.

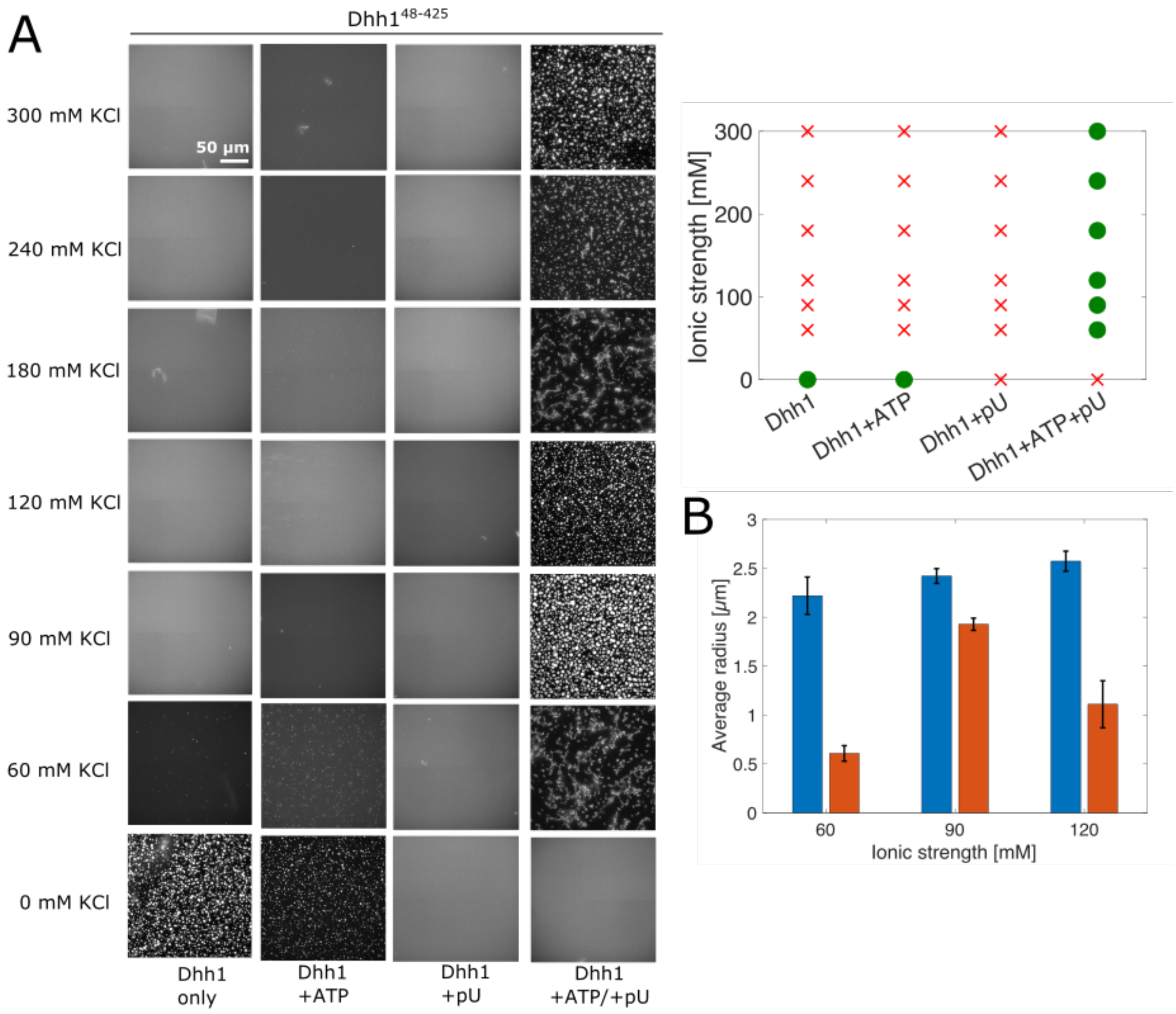


**Figure S3:** Phase diagram of Dhh1$^{48-425}$, lacking the N- and C-terminal low-complexity domains in absence / presence of ATP, polyU and ATP/polyU at increasing KCl concentration. **(A)** Left: fluorescence microscopy images; right: corresponding phase diagram. Red crosses – no phase separation, green circles – phase separation, black squares – small aggregates. **(B)** Comparison of the average droplet size of full-length Dhh1 and Dhh1$^{48-425}$ at 60, 90 and 120 mM KCl. In all cases, the droplets formed in presence of Dhh1$^{48-425}$ are smaller than in presence of full-length Dhh1

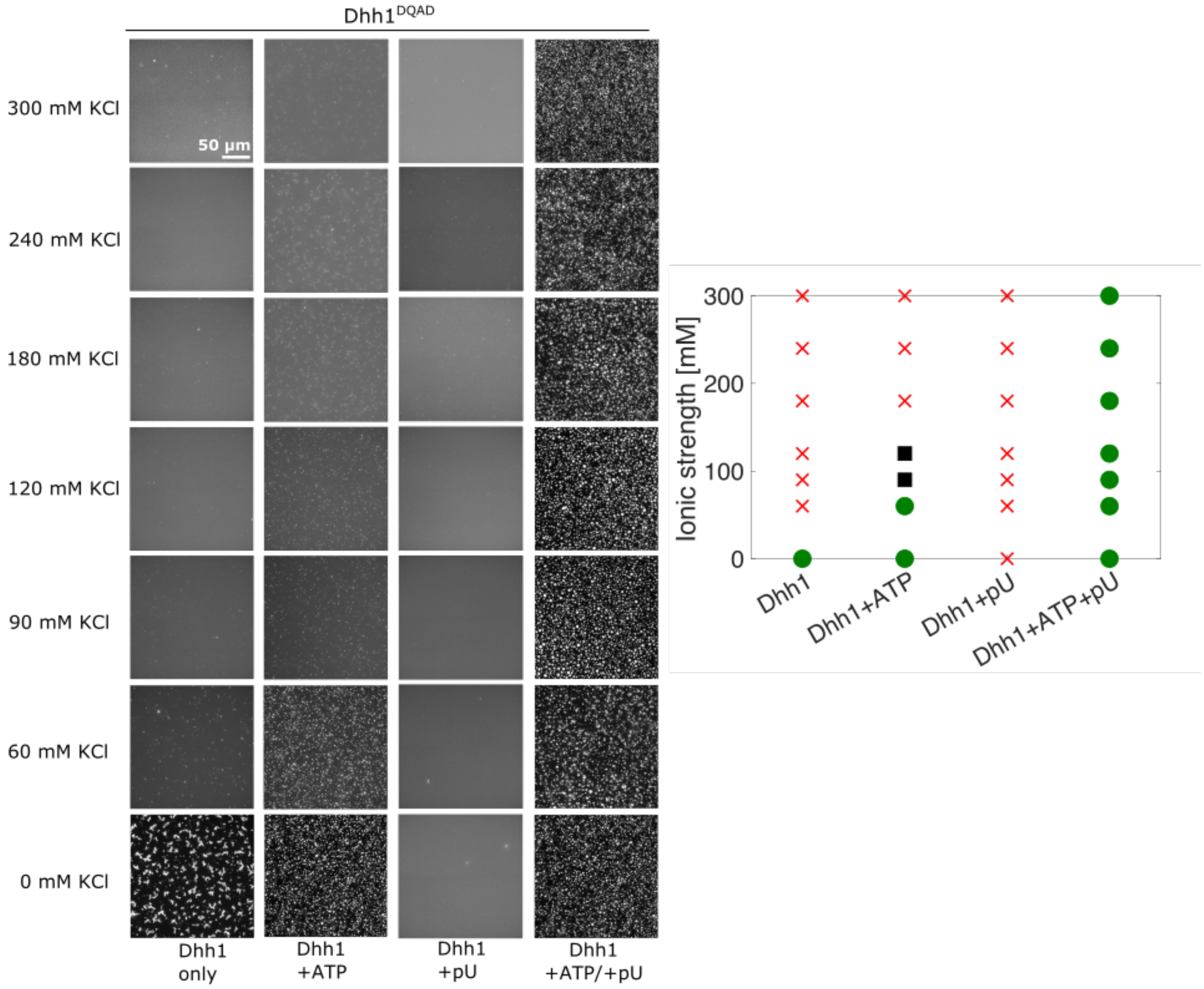


**Figure S4:** Phase diagram of Dhh1$^{DQAD}$ deficient in ATP hydrolysis propensity in absence / presence of ATP, polyU and ATP/polyU at increasing KCl concentrations. Left: fluorescence microscopy images; right: corresponding phase diagram. Red crosses – no phase separation, green circles – phase separation, black squares – small aggregates.

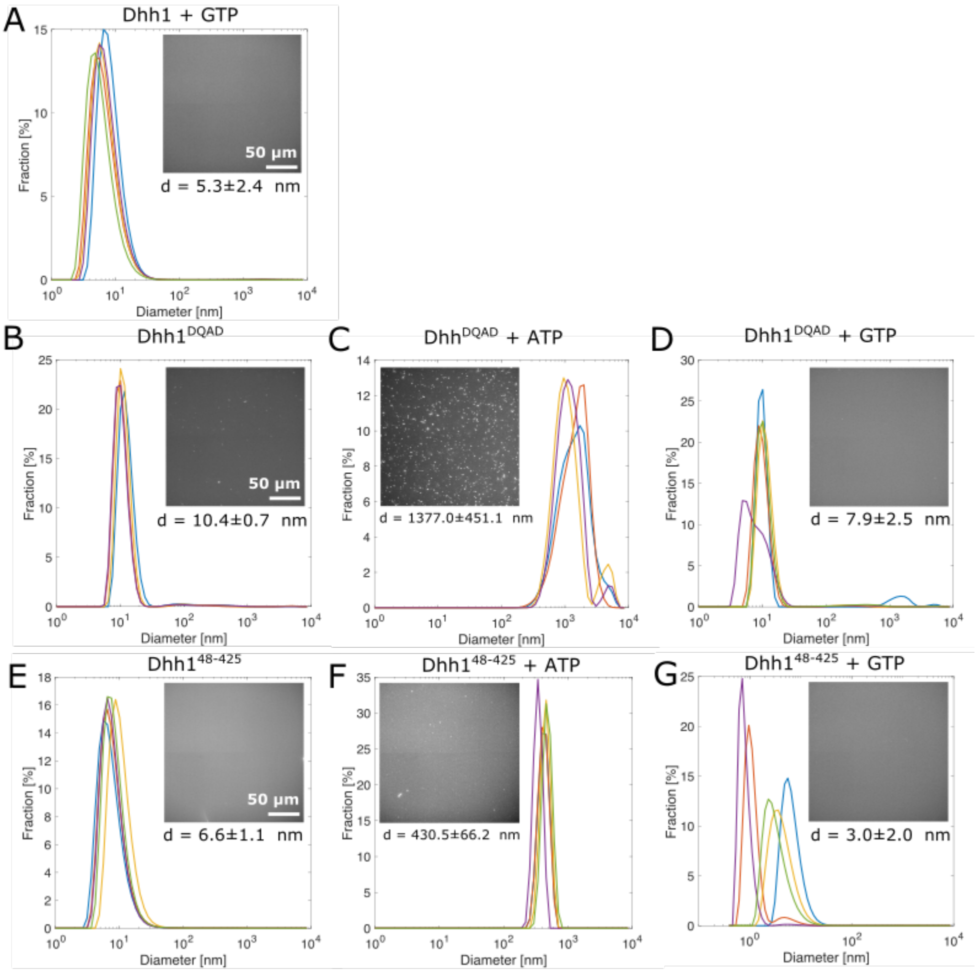


**Figure S5:** Effect of ATP and GTP on the phase transition propensity of full-length Dhh1, Dhh1$^{48\text{-}425}$, Dhh1$^{DQAD}$ measured by dynamic light scattering and fluorescence microscopy **(A)** Full-length Dhh1 remains in its monomeric form in presence of 25 mM GTP with an average diameter of 5.3 ± 2.4 nm. **(B-D)** Also Dhh1$^{DQAD}$ remains soluble in absence of ATP and GTP, as well as in presence of 25 mM GTP, but forms large condensates in presence of 5 mM ATP. **(E-G)** Dhh1$^{48\text{-}425}$ remains soluble in absence of ATP and GTP (A) and in presence of 25 mM GTP (C). Only the addition of 5 mM ATP induced the formation of small condensates with an average diameter of 430.5 ± 66.2 nm.

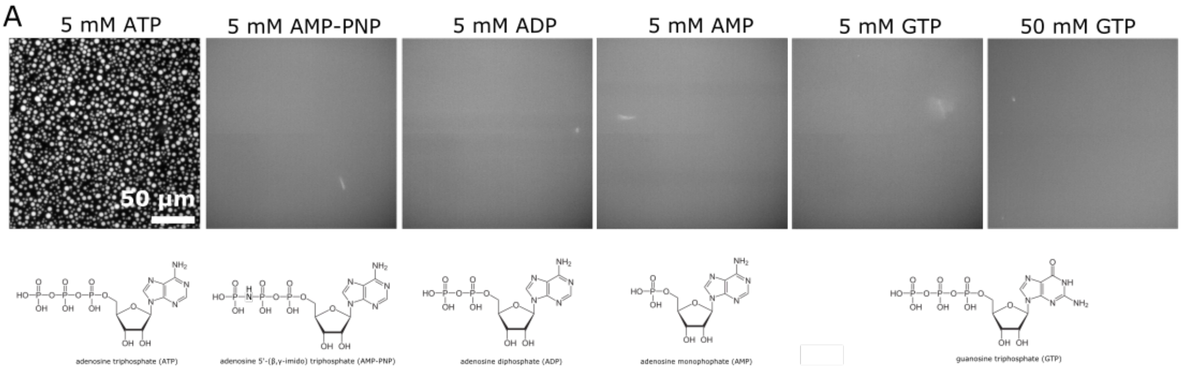


**Figure S6:** Specificity of various nucleotides to induce phase separation. Only in presence of 5 mM ATP droplet formation was induced. In presence of 5 mM non-hydrolysable AMP-PNP, ADP, AMP and GTP the full-length Dhh1 solution (including 0.5 mg/ml polyU) remained soluble. Even the addition of 50 mM GTP did not induce droplet formation, excluding effects of the ionic strength.

**Differential Dynamic Microscopy (DDM) principles and analysis.**

Within a microscopy image, the intensity value corresponding to a pixel located in position $(x, y)$ at time $t$ is denoted with $I(x, y; t)$. Fluctuations in the values of $I(x, y; t)$ can be induced by the motion of colloidal particles employed as tracers, or alternatively –in the absence of tracers– by fluctuations in the intrinsic refractive index of the sample caused by rearrangements of some structural sub-regions. A spatial variance of the type:

$$\sigma^2(\Delta t) = \sum_x \sum_y |D(x, y; \Delta t)|^2 = \sum_x \sum_y |I(x, y; t + \Delta t) - I(x, y; t)|^2$$

can thus be defined for images that are Δt apart in time. The time-averaged Fourier transform of $D(x, y; \Delta t)$, denoted with $\mathfrak{D}(\boldsymbol{q}; \Delta t) = \langle \mathcal{F}\{D(x, y; \Delta t)\}\rangle_t$, can be further radially averaged for isotropic samples, leading to the loss of the dependence on the orientation of the wavevector $\boldsymbol{q}$ and yielding a simpler $\mathfrak{D}(q; \Delta t)$ [1,2]. It can be shown that the relation:

$$\mathfrak{D}(q; \Delta t) = A(q)[1 - f(q, \Delta t)] + B(q)$$

holds, where $A(q)$ is related to the Fourier transform of the microscope Point Spread Function, $B(q)$ accounts for the camera noise, and $f(q, \Delta t)$ is the familiar Intermediate Scattering Function (ISF) of the system that can be traditionally determined by means of DLS measurements.

If an explicit analytical model for the ISF is known, as in the case of tracers that undergo Brownian diffusion in a Newtonian fluid, a straightforward determination of the rheological properties of the sample is possible by direct fitting of $\mathfrak{D}(q; \Delta t)$ [1–3].

The main advantage of DDM is represented by the fact that, unlike DLS techniques, standard white incoherent light can be used to gain structural and dynamic information in the reciprocal space starting from real-space images. Moreover, unlike standard particle-tracking rheology techniques, the tracers employed can be well below the resolution power of the microscope and do not need to be individually distinguished and tracked, leading to a robust analysis protocol free of arbitrary, user-based calibration steps.

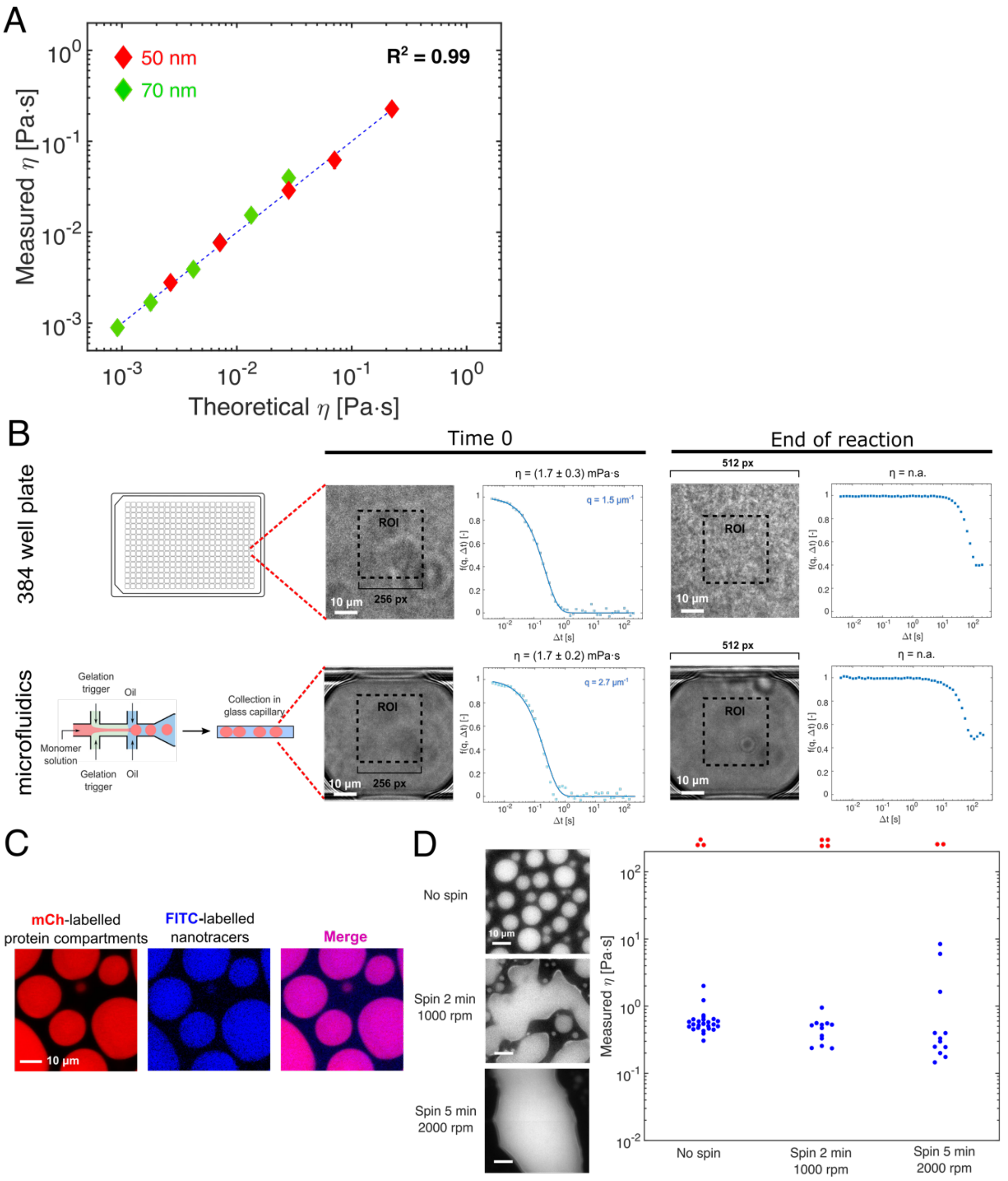


**Figure S7: Validation and control experiments for the DDM technology. (A, B)** DDM microrheology for the measurement of the viscosity of different water-glycerol mixtures with standard nanotracers (d = 50 nm, red; d = 70 nm, green) (A). From the measured diffusion coefficients *D* of the nanotracers measured by DDM, the corresponding viscosities were computed by the Stokes-Einstein equation $\eta = k_B T / 6\pi D r$, where $k_B$ is the Boltzmann constant, *T* is the absolute temperature, and *r* is the radius of the nanotracers (**B**). Error bars are smaller than the symbols. **(C)** Absence of confinement effects on the diffusion of DDM nanotracers. The gelation of a polydimethylacrylamide (PDMA) hydrogel was monitored in bulk (384-well plate) and confined in microfluidic water-in-oil droplets) to mimic phase-separated condensates. Both at time 0 and at the end of the reaction, the ISFs were comparable, showing liquid-like and arrested states, respectively, independent of the size of the container in which it was carried out. **(D)** Internalization of 25 nm nanotracers into protein-rich droplets by confocal microscopy. The fluorescence of mCherry-tagged Dhh1 and FITC-coupled nanotracers overlap perfectly. **(E)** Absence of confinement effects in condensates formed by Dhh1$^{DQAD}$. We generated compartments of increasing size by promoting the coalescence of Dhh1$^{DQAD}$ condensates by centrifugation for either 2 minutes at 1000 rpm, or 5 minutes at 2000 rpm. Despite the evident differences in morphology and size (micrographs on the *left*), the measured ISFs are consistent (panels on the *right*).

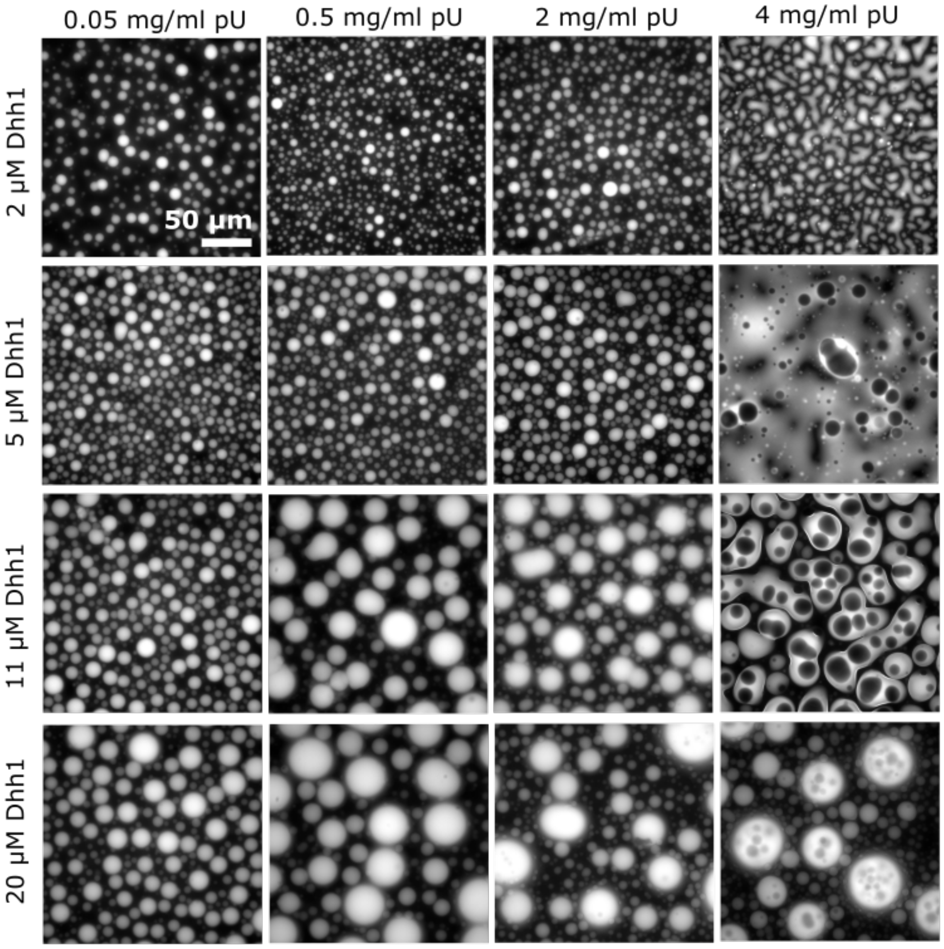


**Figure S8:** Fluorescence microscopy images, taken at various ratios of Dhh1 to polyU. At high polyU concentrations, we observed vacuole formation, most likely corresponding to a re-entrant phase transition.

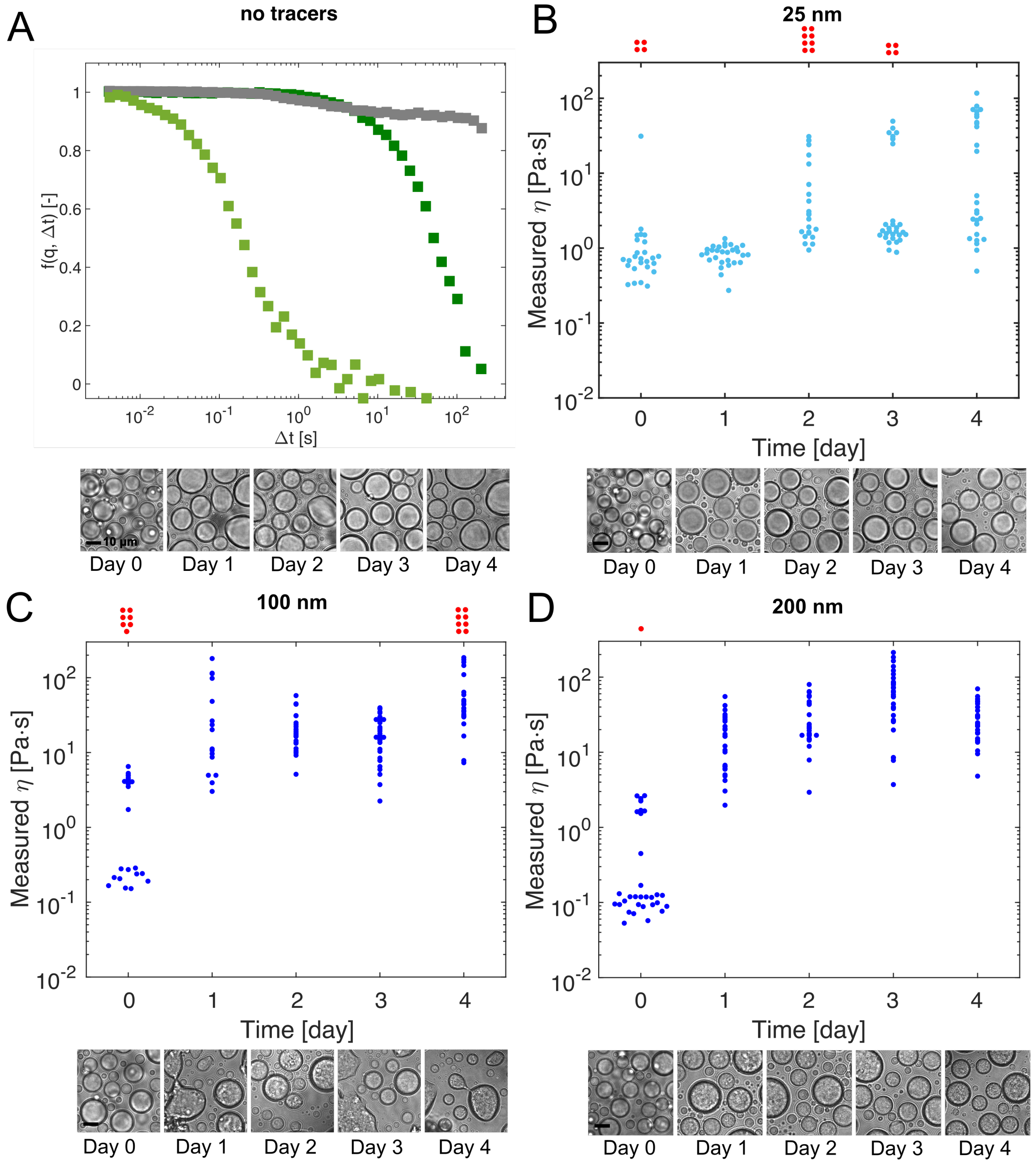


**Figure S9:** Influence of the nanotracers size on DDM measurements and compartments ageing. The ageing of phase-separated Dhh1$^{DQAD}$ compartments was investigated over 5 days in the absence of tracers (**A**) and in the presence of nanotracers with diameter of 25 nm (**B**), 100 nm (**C**), or 200 nm (**D**). When no tracers are added to the phase-separating mixture (**A**), a non-null DDM signal is provided, most likely originating from intrinsic fluctuations in the refractive index of structural sub-domains within the compartments. The characteristic size of these domains is unknown, and may even vary within the droplet. This hinders the computation of absolute viscosity values, and accounts for more dispersed estimates than those observed when using nanotracers of definite size and reduced polydispersity. Interestingly, however, the DDM measures performed without nanotracers qualitatively confirmed the fractions of liquid, low and high viscous, as well as dynamically arrested states on 4 consecutive days, observed when using the standard 25 nm tracers (**B**). Therefore, we can exclude artefacts induced by the presence of the nanotracers and/or by their interactions with the components of the compartments. Tracers of 100 and 200 nm (**C**, **D**) were observed by bright field microscopy to dramatically interfere with the structure and ageing of the compartments (**E**), yielding results that significantly differ from those obtained with smaller/no tracers and making them unsuitable for reliable DDM analysis. Scale bars are 10 μm.

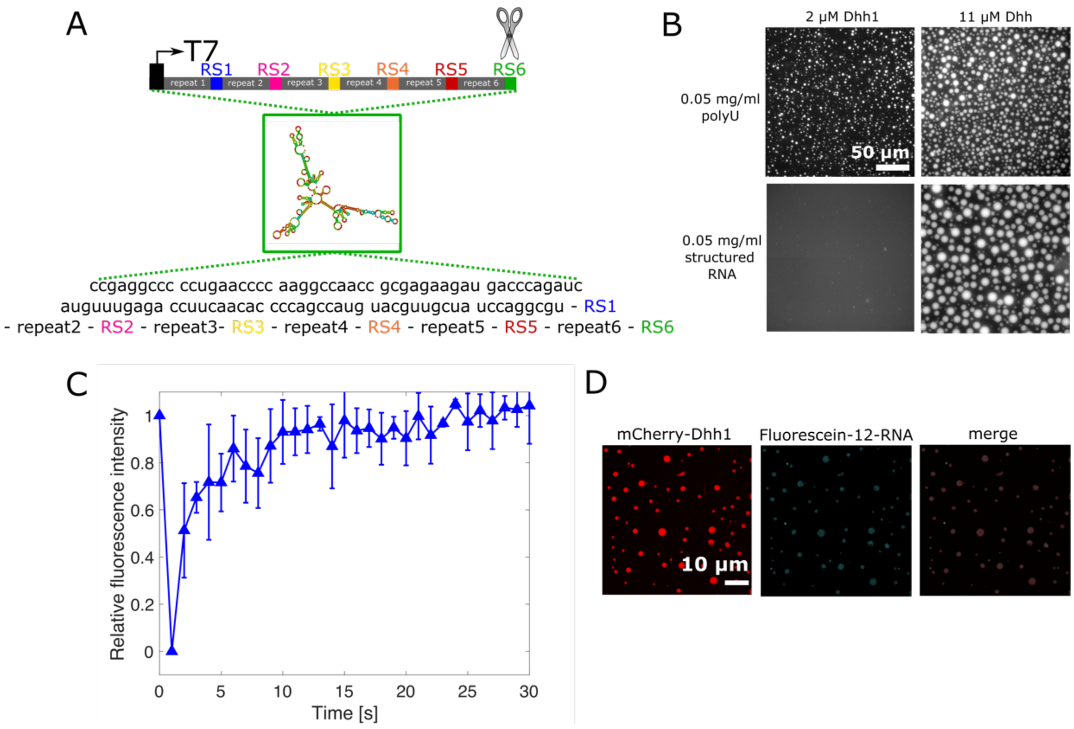


**Figure S10:** Effect of structured RNA on the phase separation propensity and material properties of Dhh1 condensates. **(A)** In vitro transcribed construct of repetitive 100 nucleotides of the actin mRNA sequence (sequence source) linked by six different restriction sites for six different restriction enzymes. To obtain the 600 nt RNA used in this study, the plasmid (pET-15b) was linearized by digestion with BamHI (RS6) and in vitro transcribed by T7 RNA polymerase. **(B)** Droplet formation in presence of unstructured polyU and structured RNA. In presence of 0.05 mg/ml polyU, droplets could be formed in presence of 2 µM Dhh1 and 5 mM ATP, whereas in presence of 0.05 mg/ml structured RNA 11 µM Dhh1 were required**.** **(C)** Fluorescence recovery after photobleaching (FRAP) of Dhh1 droplets formed in presence of 0.05 mg/ml polyU and 11 µM Dhh1 after 2 h of incubation shows that these droplets fully recover and thereby remain highly dynamic over the observed time scale. **(D)** Droplet heterogeneity, shown by varying single droplet fluorescence intensity of mCherry-tagged Dhh1 and Fluorescein-12-labeled 600 nt structured RNA.
